# Molecular recognition of sugar binding in a melibiose transporter MelB by X-ray crystallography

**DOI:** 10.1101/2020.12.20.423627

**Authors:** Lan Guan, Parameswaran Hariharan

## Abstract

The symporter melibiose permease MelB is the best-studied representative from MFS_2 family and the only protein in this large family with crystal structure determined. Previous thermodynamic studies show that MelB utilizes a cooperative binding as the core mechanism for its obligatory symport. Here we present two sugar-bound X-ray crystal structures of a *Salmonella typhimurium* MelB D59C uniport mutant that binds and catalyzes melibiose transport uncoupled to either cation, as determined by biochemical and biophysical characterizations. The two structures with bound nitrophenyl-α-D-galactoside or dodecyl-β-D-melibioside, which were refined to a resolution of 3.05 or 3.15 Å, respectively, are virtually identical at an outward-facing conformation; each one contains a α-galactoside molecule in the middle of protein. In the substrate-binding site, the galactosyl moiety on both ligands are at an essentially same configuration, so a galactoside specificity determinant pocket can be recognized, and hence the molecular recognition mechanism for the binding of sugar in MelB is deciphered. The data also allow to assign the conserved cation-binding pocket, which is directly connected to the sugar specificity determinant pocket. The intimate connection between the two selection sites lays the structural basis for the cooperative binding and coupled transport. This key structural finding answered the long-standing question on the substrate binding for the Na^+^-coupled MFS family of transporters.

**Significance:** Major facilitator superfamily_2 transporters contain >10,000 members that are widely expressed from bacteria to mammalian cells, and catalyze uptake of varied nutrients from sugars to phospholipids. While several crystal structures with bound sugar for other MFS permeases have been determined, they are either uniporters or symporters coupled solely to H^+^. MelB catalyzes melibiose symport with either Na^+^, Li^+^, or H^+^, a prototype for Na^+^-coupled MFS transporters, but its sugar recognition has been a long-unsolved puzzle. Two high-resolution crystal structures presented here clearly reveal the molecular recognition mechanism for the binding of sugar in MelB. The substrate-binding site is characterized with a small specificity groove adjoining a large nonspecific cavity, which could offer a potential for future exploration of active transporters for drug delivery.

## Main

Membrane transporters play critical roles in human health and disease, and they are potential drug targets with increasing interests in pharmaceutical applications. While active transporters can move specific solutes into cells effectively, a majority of available drugs enter cells by an inefficient solubility diffusion. This outcome was largely due to lack of knowledge on active transporters, in particular the detailed understanding about substrate-binding sites and transport mechanisms. The glycoside-pentoside-hexuronide:cation symporter family (GPH)^(1)^, which is a subgroup of the major facilitator superfamily (MFS) of membrane transporters found from bacteria to mammalian cells that catalyze a coupled substrate transport with monovalent Na^+^, Li^+^, or H^+^, Na^+^ or Li^+^, H^+^ or Li^+^, or only H^+(2)^. Most members transport glycosides, but the substrates transported by this family are diverse from small molecules, such as glucose, melibiose, raffinose, or sulfoquinovose, to bigger molecules such as phospholipids. For example, the Na^+^-coupled MFSD2A in brain and eyes that mediates the delivery of the essential omega-3 fatty acids for neurodevelopment^(3)^. The human MFSD13a transporter, which is expressed in many tissues but overexpressed in colorectal cancer cells, has been successfully targeted for diagnosis and treatment^(4, 5)^. Based on the larger Pfam database, these proteins belong to MFS_2 family^(6)^ with greater than 11,000 sequenced genes across thousands species (**https://pfam.xfam.org/family/PF13347#tabview=tab0**). So far, there is only one member with high-resolution crystal structure published^(7)^. Lack of high-resolution substrate-bound 3-D structure from this large family of transporters is a bottleneck for unraveling molecular recognition and transport mechanisms, which significantly hampered the developments towards the pharmaceutical applications as drug targets or as vehicles for drug delivery.

The bacterial melibiose permease MelB is one of the few membrane transporters discovered at earlier years^(8)^, and it has been serving as a model system for the study of cation-coupled transport mechanisms^(9-12)^, as well as the development of novel analytical techniques for membrane protein research^(11, 13-17)^. It is a well-documented representative from MFS_2 family^(1, 7, 10, 12, 18-22)^. MelB catalyzes the stoichiometric galactose or galactosides symport with monovalent cation either Na^+^, Li^+^, or H^+^, with no affinity for glucose or glucosides^(23-26)^. The symport reactions are reversible, and the polarity of transport is determined by the net electrochemical gradients from both substrates. In either symport process, the coupling between the driving cation and driven sugar is obligatory^(9, 25, 27, 28)^. The previous X-ray 3-D crystal structure of the MelB of *Salmonella typhimurium* (MelB_St_) shows a MFS fold at an outward-facing conformation^(7)^; while no bound sugar or cation was resolved, this structure localizes the residues that have been functionally determined to be important for the co-substrate binding within a large cavity^(7, 9, 29-36)^. Like other MFS transporters^(37-42)^}, the well-recognized alternating-access mechanism has been also proposed in MelB^(21, 29, 34)^.

The alternating-access mechanism only describes how the conformational changes to create the substrate-access path alternatively at both surface of transporters during transport; however, how the proteins respond to its specific substrate binding and initiate the alternating-access mechanism for transport are generally not so clear. Recently, with MelB_St_, the free energy for the binding of Na^+^ and melibiose were determined with the isothermal titration calorimetry (ITC)^(22)^. A binding thermodynamic cycle indicated that a core mechanism for this symporter is positive cooperativity of the co-substrate (melibiose vs. Na^+^ or H^+^)^(22)^. To further gain the structural basis, we determined two galactoside-bound crystal structures of an uniport mutant D59C MelB_St_ with a mutation at a highly conserved cation site, which enabled deciphering the molecular recognition mechanism for the binding of sugar in MelB.

Asp59 on the helix II has been well-studied with MelB of *Escherichia coli* (MelB_Ec_)^(34, 43, 44)^. The D59C mutation in both MelB_Ec_ and MelB_St_ abolishes Na^+^ binding and Na^+^ stimulation on galactoside binding or transport, but catalyzes melibiose transport independent of the cation^(7, 22, 34, 43, 44)^. It has been determined that Asp59 is the ligand for Na^+^ and also for H^+^; the D59C mutant behaves like a uniporter. Two crystal structures of D59C MelB_St_ with a bound melibiose analogue 4-nitrophenyl-α-D-galactopyranoside (α-NPG) or a novel melibiose-derived detergent n-dodecyl-β-melibioside (DDMB) were determined. Both structures are essentially identical; all exhibits an open periplasmic vestibule accessing to the bound sugar without bound cation, representing the binary complex of MelB with bound single galactopyranoside at an outward-facing conformation.

## Results

### Melibiose transport and ligand binding

[^3^H]melibiose transport assays confirmed that the D59C MelB_St_ mutant fails in catalyzing all three modes of melibiose active transport coupled to H^+^, Na^+^, or Li^+(7)^, even with increased concentration of Na^+^ or Li^+^ (**Fig. 1a**); however, it supports a normal melibiose fermentation on MacConkey agar plates containing melibiose as the sole carbon source (**Fig. 1a insets**). This melibiose transport rate-dependent color development assay indicates that melibiose translocation mechanism is retained with this mutant.

**Fig. 1.**
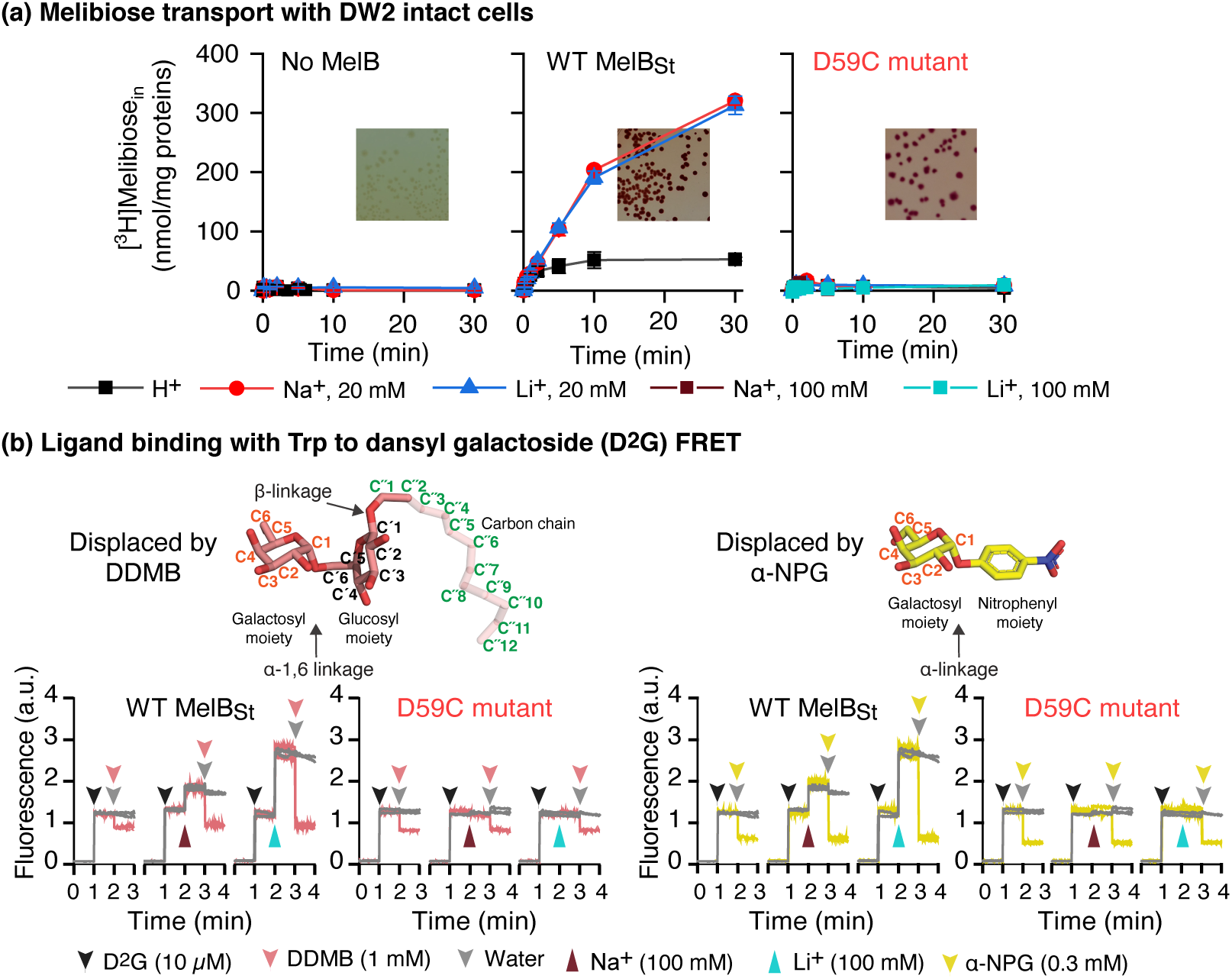
Functional characterizations. **(a)** [^3^H]Melibiose transport time course with intact cells. *E. coli* DW2 strain (*melB, lacZY*) expressed with WT MelB_St_ or D59C MelB_St_ mutant were subjected to [^3^H]melibiose active transport assay at 0.4 mM and 10 mCi/mmol in the presence of 20 mM or 100 mM NaCl or LiCl by a fast filtration method. Inset, melibiose fermentation assay. Transformants with DW2 strain were plated on MacCokey agar plates containing 30 mM melibiose as the sole carbon source and pH sensor neutral red, and incubated at 37 °C for 16-18 hr before imaging. **(b)** Trp→D^2^G FRET. Purified proteins at 1 μM concentration in 20 mM Tris-HCl, pH 7.5, 100 mM CholCl, 10% glycerol, and 0.03% UDM were subjected to steady-state fluorescence measurements as described in Methods. On the time-trace, D^2^G at 10 μM was added into the protein solutions at 1-min point, followed by supplying DDMB or α-NPG at a saturating concentration to displace the bound D^2^G. For testing Na^+^ or Li^+^ binding, they are added into the solution before displacement with the second sugar. The sugar moiety on DDMB or α-NPG is shown in chair form and carbon positions are labeled. Glycosidic bonds are indicated by arrows. Both sugars are α anomers of galactosides containing a substituent at the anomeric carbon-1 at an α form.

ITC measurements were used to analyze the co-substrate binding **(Supporting Information Fig. S1; Table 1)**. The cooperative binding between Na^+^ and melibiose with WT MelB_St_ has been reported ^(22)^. The D59C mutant at apo state exhibits a *K*_d_ value for melibiose with a surprisingly 2-fold better than the WT; however, no Na^+^ or Li^+^ effect on melibiose binding was detected. Consistently, α-NPG also binds to apo D59C mutant at an approximately 4-fold increased affinity, with no Na^+^ or Li^+^ effect. Li^+^ binding measurements to the WT show a *K*_d_ value at apo state of 0.40 ± 0.10 mM, which decreased by 6-fold in the presence of melibiose; notably, the cooperativity is less than melibiose with Na^+^ with a 8-fold change^(22)^. For D59C mutant, all heat changes derived from Na^+^ or Li^+^ binding are quite small even with increased concentration or in the presence of melibiose, which prevents an accurate determination of the dramatically increased *K*_d_. Collectively, D59C MelB_St_ selectively eliminates the capabilities in binding of cation and its role in cooperating with melibiose, as well as their co-transport, but the capability for melibiose binding and translocation is retained; thus, D59C MelB_St_ mutant is an uncoupled mutant as proposed for MelB_Ec_ ^(22)^.

**Table 1.** *K*_d_ of substrate binding to MelB_St_

| | Conditions | Melibiose (mM) | α-NPG (μM) | Conditions | Na<sup>+</sup> binding (mM) | Li<sup>+</sup> binding (mM) |
| --- | --- | --- | --- | --- | --- | --- |
| WT | Apo | 9.28 ± 0.23<sup>*†</sup><br>(n = 3) | 43.20 ± 0.62<br>(n = 2) | Apo | 0.68 ± 0.03<sup>†</sup><br>(n = 4) | 0.40 ± 0.10<br>(n = 2) |
| | Na<sup>+</sup> (100 mM) | 1.09 ± 0.06<sup>†</sup><br>(n = 4) | 25.69 ± 4.42<br>(n = 2) | Melibiose (50 mM) | 0.08 ± 0.01<sup>†</sup><br>(n = 4) | 0.07 ± 0.01<br>(n = 2) |
| D59C | Apo | 4.66 ± 0.10<br>(n = 3) | 10.47 ± 1.34<br>(n = 2) | Apo | /<sup>#</sup><br>(n=2) | /<br>(n=2) |
| | Na<sup>+</sup> (100 mM) | 4.96 ± 0.11<br>(n = 3) | 11.97 ± 0.09<br>(n = 2) | Melibiose (50 mM) | /<br>(n=1) | /<br>(n=1) |

<sup>\*</sup>SE, Standard error; <sup>#</sup>, curve fitting was not carried out; n, the number of tests; <sup>†</sup>, data from the publication in reference 22 or averaged with more tests.

DDMB is a melibiose-derived novel detergent containing a melibiosyl moiety and a 12-carbon chain^(45)^ **(Fig. 1c)**. A well-established FRET measurement^(26, 33)^, which is based on MelB Trp residues to dansyl moiety on the bound fluorescent analogue dansyl-2-galactoside (D^2^G), was used to determine MelB_St_ affinity for DDMB. Addition of DDMB or α-NPG into the D^2^G-bound WT MelB_St_ solution leads to a large decrease in the fluorescent intensity as a result from the exchange of bound D^2^G by MelB_St_ with DDMB **(Fig. 1b, pink)** or α-NPG **(Fig. 1b, yellow)**, strongly indicating that DDMB can competitively bind to the WT MelB_St_. Na^+^ or Li^+^ stimulation on the FRET with WT is typically shown as the increase in fluorescent intensity upon adding NaCl or LiCl (at 2-min time point) prior to the displacement. The data clearly show that positive cooperativity with Na^+^ or Li^+^ also exists with DDMB. D59C mutant also binds DDMB with no Na^+^ or Li^+^ stimulation as expected.

Quantitatively, the half-maximal concentration for displacing the bound D^2^G (IC_50_) was determined for DDMB and α-NPG (**SI Fig. S2**). Titration of DDMB was carried out at a concentration less than CMC value (0.295 mM) to keep DDMB in monomeric form, and the FRET intensity was decreased along with the increase in DDMB concentrations. As a negative control, the maltoside-based detergents DDM and UDM were tested, which only shows a dilution effect similar to the addition of water and supports the specific effect observed from DDMB. The curve fitting reveals IC_50_ value of 26.13 ± 5.46 μM with the apo WT MelB_St_, which is decreased by 5-fold to 4.79 ± 0.28 μM in the presence of Na^+^ **(Table 2; SI Fig. S2)**. Since both monomer and micelles co-exist in the concentrated solutions used for the titration, the IC_50_ values might be underestimated. Even so, DDMB is the highest affinity ligand among all tested galactosides to MelB_St_. The D59C mutant yields a IC_50_ of 20 μM for DDMB; consistently, there is little effect by Na^+^. Use of this high-affinity substrate further confirms the uncoupling property with this mutant.

**Table 2.** IC_50_ of galactoside displacement on bound D^2^G by MelB_St_

| | DDMB (μM) | | α-NPG (μM) | |
| --- | --- | --- | --- | --- |
|  | Apo | Na<sup>+</sup> (100 mM) | Apo | Na<sup>+</sup> (100 mM) |
| WT | 26.13 ± 5.46<br>(n = 3) | 4.79 ± 0.28<br>(n = 3) | 58.17 ± 11.53<sup>*</sup><br>(n = 2) | 22.96 ± 2.24<br>(n = 3) |
| D59C | 20.16 ± 7.37<br>(n = 2) | 23.56 ± 3.90<br>(n = 2) | 22.15 ± 5.75<br>(n = 2) | 21.27 ± 1.67<br>(n = 2) |

<sup>\*</sup>SE, standard error. n, the number of tests.

### X-ray crystal structures of D59C MelB_St_ with bound galactoside

Purified D59C MelB_St_ in UDM solution exhibits an improved thermostability with a melting temperature (*T*_m_) of 60 **°**C as detected by circular dichroism (CD) spectroscopy, which is greater by 6 **°**C than the WT (**SI Fig. S3**)^(46)^. Both proteins are well-folded with α-helical secondary structure overwhelmingly dominated. Two crystal structures of D59C MelB_St_ with bound α-NPG or DDMB were modeled from positions 2 to 454 without a gap (**Fig. 2a**), refined to a resolution of 3.05 Å and 3.15 Å, and assigned with a PDB access number of 7L17 and 7L16, respectively. The phase was determined by MR-SAD based on selenomethionine (SeMet)-incorporated D59C MelB_St_ and a model from structure pdb id, 4m64^(7)^ (**SI Fig. A4a, b)**. The two native structures are superimposed very well, with rmsd of 0.187 Å, showing a typical MFS fold (**Fig. 2a**) with 12 transmembrane helices as described. The cytoplasmic middle loop carrying a short α-helix links the two helical bundles formed by the N- and C-terminal six α-helices; the C-terminal tail also contains a short α-helix with the last 26 residues unassigned due to poor electron densities.

**Fig. 2.**
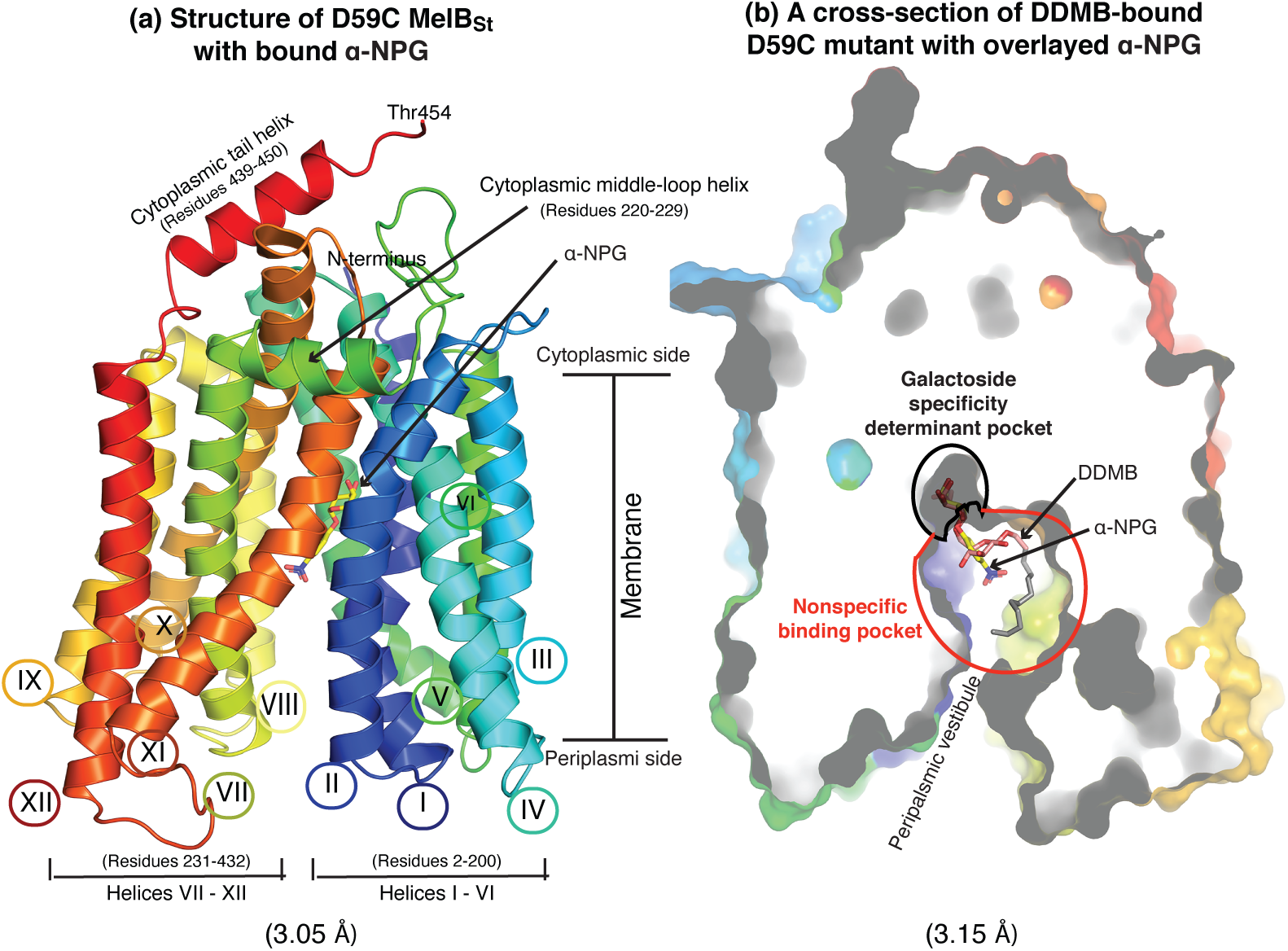
X-ray crystal structures of D59C MelB_St_ bound with α-NPG or DDMB. **(a) Overall fold and helical packing**. The D59C MelB_St_ structure bound with α-NPG is viewed parallel to the membrane and shown in cartoon representation, which is colored in rainbow from N-terminus blue to C-terminal end in red. Both termini are located in the cytoplasmic side as indicated. The membrane-spanning region is proximately estimated and indicated. Transmembrane helix is numbered in Roman numerals, and two cytoplasmic helices is named as cytoplasmic middle-loop helix or cytoplasmic tail helix. The amino-acid sequence positions for the N-terminal 6 helices (I-VI) and the C-terminal 6 helices (VII-XII), as well as the two cytoplasmic helices, are indicated. The bound α-NPG is indicated. **(b) Outward-facing conformation bound with DDMB**. The structure is shown in a cross-section of surface representation with the N-terminal bundle orientated at the left side. DDMB is shown in pink and tail (C2-12) in gray. The galactoside specificity determinant pocket is indicated by a black cycle, and the non-specific cavity is indicated by a red cycle. A α-NPG molecule was included that was from the α-NPG-bound structure after overlayed with the DDMB-bound structure. PBD identification code for both structures are indicated.

An open cavity at periplasmic side is formed by 8 inner-layer helices (I, II, IV, V, VII, VIII, X, and XI) (**SI Fig. S5**); at the apex of the cavity in both structures, one molecule of α-NPG (**Fig. 2a)** or DDMB (**Fig. 2b)** is bound to the sugar-binding pocket as indicated. This pocket is connected to the periplasmic surface via the solvent-accessible periplasmic vestibule that allows the sugar to access to the sugar-binding site. Overlay of the two structures shows a narrow pocket that only hosts the specific galactosyl moiety of both ligands as indicated as black cycle in panel b, named galactoside specificity determinant pocket. The cavity as indicated by the red cycle big enough to accommodate a detergent tail with 12 carbons, and named as non-specific cavity. More details will be discussed in the followed sections. There is no solvent-accessible tunnel towards the cytoplasmic surface. The cytoplasmic tail together with helices IV-loop-V, X-loop-XI, as well as middle loop, stabilize the inside-closed, outside-open binary complex of MelB_St_ with bound galactoside.

### α-NPG binding site

As shown in **Fig. 3a**, a strong electron density blob links the N-terminal helices I and IV, which fits well to a α-NPG molecule that was presented in the crystallization. The galactosyl moiety lies in a small pocket forming multiple close contacts with the N-terminal helices (I, IV, and V) and few interactions with the C-terminal helices (X and XI) (**Fig. 3b**). The substituent nitrophenyl group is exposed to the large solvent-filled cavity, with much less polar interactions with the protein. In the cavity, there are some unassigned densities that also presents in the DDMB-bound structure. The molecular recognition of α-NPG by MelB are highly focused around the specific galactosyl moiety. Each of the four hydroxyl groups (OH) are fully liganded by 2-3 hydrogen-bonds (H-bond, mainly bifurcated H-bonds), and the galactosyl ring is sandwiched by aromatic sidechains vie Ch-π interactions. The two negatively charged Asp19 (helix I) and Asp124 (helix IV) make the major contributions to the four OH (**Fig. 3a, b**). Asp19 and Asp124 accept hydrogen atoms (H) from C2- and C3-OH, or C4-OH and C6-OH of the galactosyl moiety, at distances less than 3.5 Å. Asp124, at a H-bond distance to Trp128, also forms a salt-bridge interaction with Lys18 (helix I), which supports Lys18 to donate a H to C3-OH by stabilizing this H-bond donor at a position between the two H-bond acceptors Asp19 & 124. Furthermore, Arg149 donates two H bonds to C2-OH; Trp128 also donates a bifurcated H-bond to C3- and C4-OH. From the C-terminal bundle, Lys377 and Gln372 (helix XI) are within 5 Å distance from C6-OH. Tyr120 (helix IV) forms a H-bond with Lys18 with no close contact (>3.5 Å distance) to the sugar; The373 (helix XI) form a H-bond with Asp124, which is the only interaction from the C-terminal bundle to the galactosyl-binding site with no close contact to the sugar. Characteristically, all these interactions link around galactosyl moiety, which forms a widespread salt-bridge assisted H-bonding network to “hang” the sugar in the middle of protein **(Figs. 1, 3c)**.

**Fig. 3.**
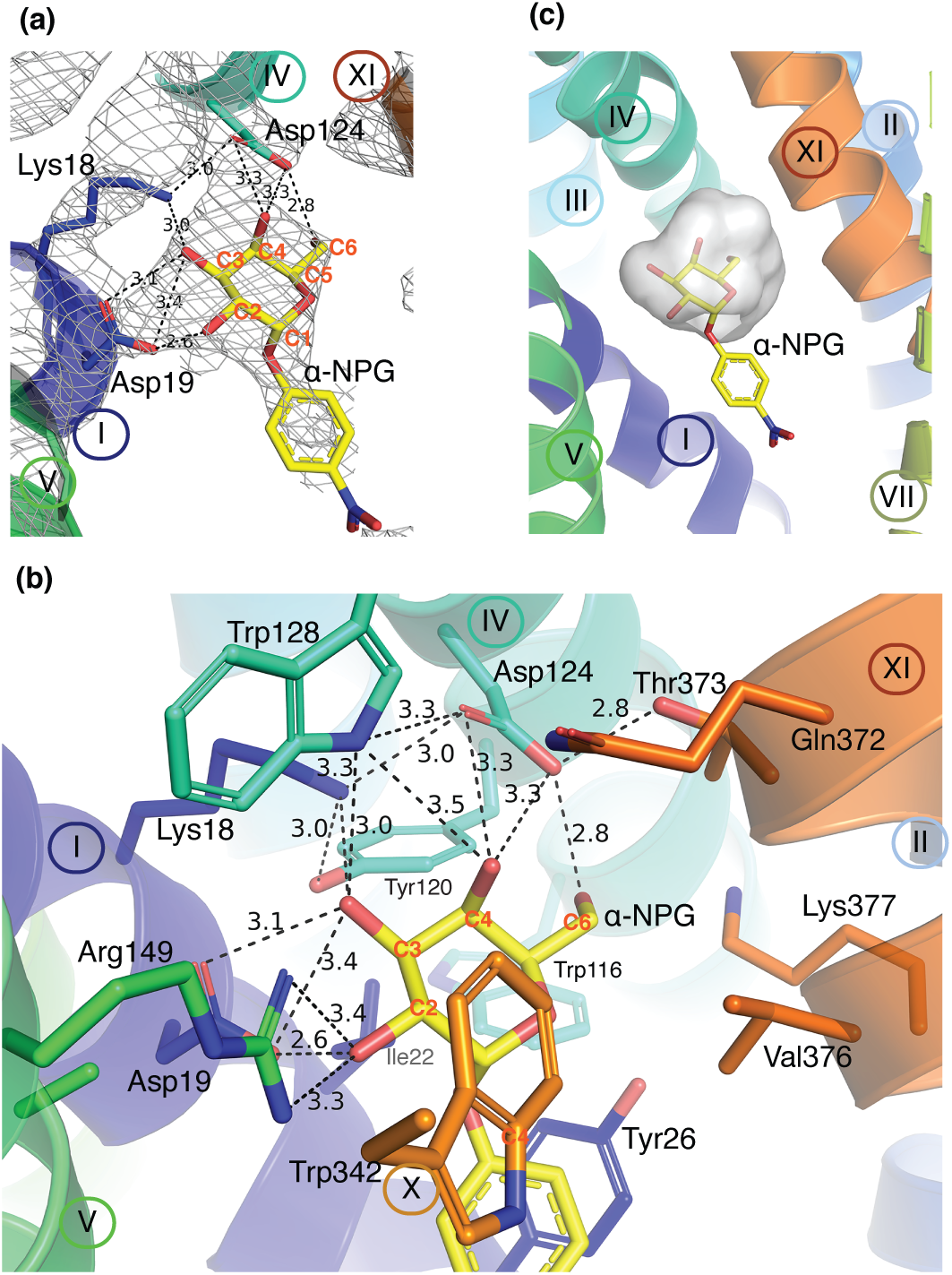
α-NPG-binding site. The structure with bound α-NPG was refined to a resolution of 3.05 Å [PDB ID, 7L17]. Helices are shown in a cartoon representation and labeled in Roman numerals; protein sidechains are shown in sticks, labeled in three letters, and color-matched with hosting helix. This style is used throughout this article. Carbon positions on the galactosyl ring are labeled in red. H-bond and salt-bridge interactions are judged by distance that is ≤ 3.5 Å and indicated by dashed lines with distance shown in Å. The α-NPG molecule is shown in stick and the carbons are colored in yellow with oxygen in red and nitrogen in blue. **(a) 2Fo-Fc electron-density map**. One molecule of α-NPG is fitted to the electron density (contoured to 1.5 σ). Residues Asp19&Lys18 (helix I) and Asp124 (Helix IV) form multiple strong H-bonds with all four OH groups on the galactosyl ring of α-NPG. **(b) Sugar-binding site**. Helix I (residues Lys18 and Asp124), helix IV (residues Tyr120, Asp124, Trp128), helix V (Ala152, Arg149), and helix XI (residues Thr373) are involved in a charge-assisted H-bond network with all four OH on the specific galactosyl ring. Ile22 and Tyr26 (helix I), Trp342 (helix X), Val376 (helix XI) on helix IV are approximately 4 Å away and provide hydrophobic environment for galactosyl ring and/or phenyl ring. Few residues at longer distances are also shown up including Trp116 (helix IV) and Gln372 & Lys377 (helix XI). **(c) Galactoside specificity determinant pocket**. The binding residues listed in panel (b) is shown in cavity presentation and colored in gray, where only hosts the galactosyl moiety of the α-NPG molecule.

Four aromatic residues are present in this sugar-binding cavity (**Fig. 3b**). Trp128 and Tyr120 were engaged in this H-bonding network. Trp342 (helix X) shapes the binding pocket by providing Ch-π interactions with both of the galactosyl and phenyl rings, and it also form aromatic stacking with Trp128. Tyr26 (helix I) forms an aromatic stacking interaction with the phenyl ring of α-NPG and indole ring of Trp116 (helix IV), and it could also form a potential H-bond with Lys377. Ile22, Trp26, Trp116, and Als152 (Helix V) lie on the relatively hydrophobic face of the galactosyl ring; Trp342 and Val376 (helix XI) lie on the other face; these residues provide a hydrophobic environment favored by the phenyl ring. In addition, Asn251, at 4.5 Å distance to the nitro group in the *para* position, together with further distancing Asn248 and Asn244 on the same face of helix VII, may contribute a polar environment for this nitro group. These chemical environments around the nitrophenyl moiety could explain why α-NPG exhibits 100-fold greater affinity compared to the native substrate melibiose (**Table 1**).

### DDMB-binding site

The DDMB-bound structure of D59C mutant is indistinguishable from that with α-NPG, but the electro-density blob in the galactoside-binding pocket differs from that for α-NPG. It fits fairly well with a melibiosyl moiety of DDMB **(Fig. 4)**. Notably, the β-glycosidic bond on C1′ position is supported by the electron-density map, and the C3-12 on the carbon chain tail exhibits weak or no electron-density with occupancy set to 0 or 0.3 and colored in gray. Overall, the galactosyl moiety is in a configuration essentially indistinguishable from that of α-NPG. The glucosyl ring on melibiosyl moiety is nearly perpendicular to the galactosyl ring, which makes a bifurcated H-bond with Trp342 on the C1′-O atom and C2′-OH group. In addition, the polar residues Asn244 (helix VII) is 4.8 Å away from C1′-O atom, and Ser153 (helix V) and Thr338 (helix X) are at distances of <6.0 Å to C3′- and C2′-OH, which contribute a polar environment around the glucosyl moiety. This configuration of melibiosyl moiety may provide clues how the native substrate melibiose binds to MelB_St_.

**Fig. 4.**
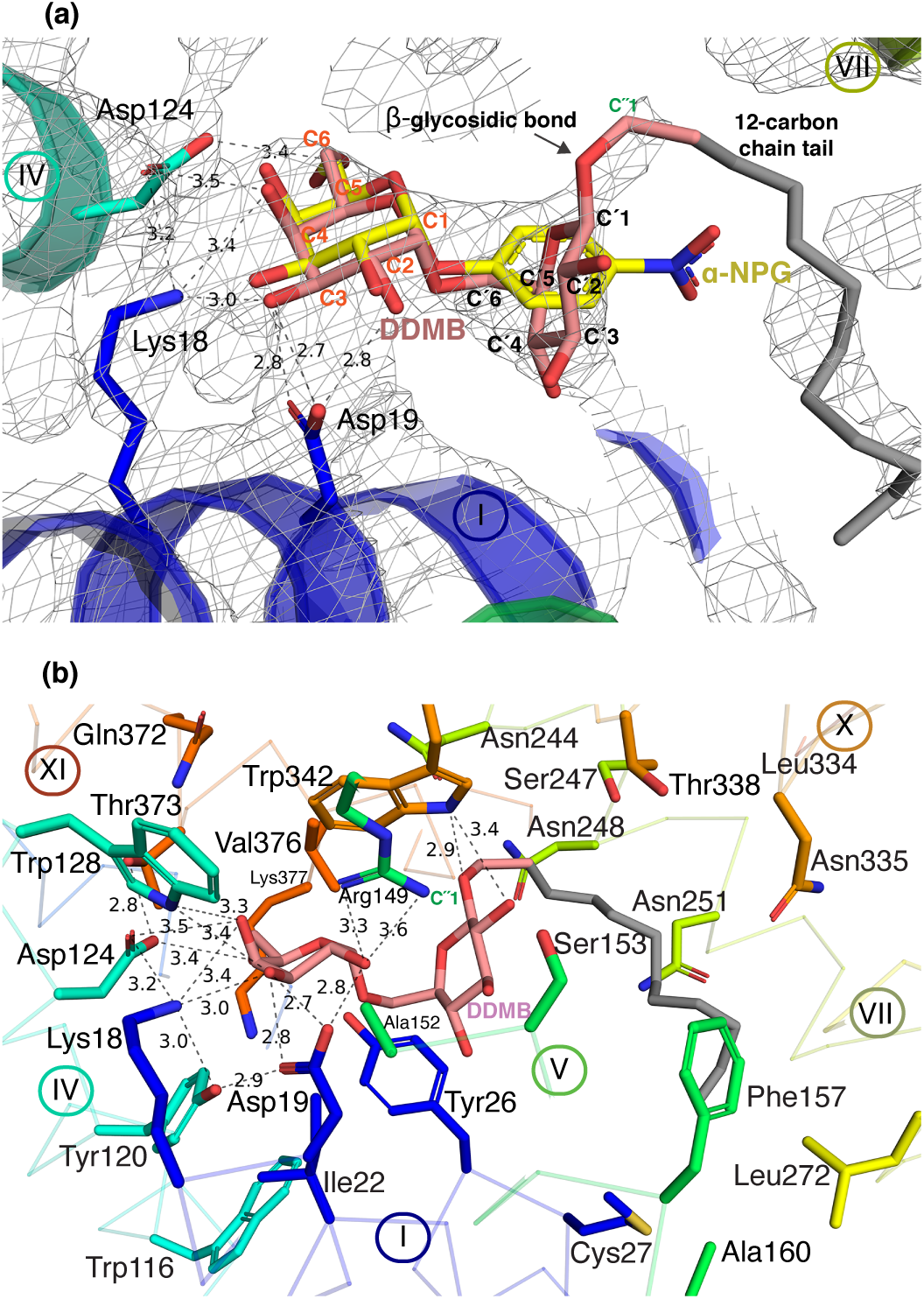
DDMB-binding site. The structure with bound DDMB was refined to a resolution of 3.15 Å [PDB ID, 7L16]. Carbon positions are indicated (also presented in **Fig. 1**). The C3 to C12 on the detergent chain tail are presented in transparency due to lack of electron density. H-bond and salt-bridge interactions are judged by distance ≤ 3.5 Å and indicated by dashed lines with distance shown in Å. **(a) 2Fo-Fc electron-density map**. One molecule of DDMB is fit to the density map (contoured to 1.0 σ). The α-NPG in the α-NPG-bound structure, which was aligned to the DDMB-bound structure, is left for comparison. **(b) DDMB-binding site**. Similar contacts between MelB_St_ and the galactosyl moiety observed in the α-NPG binding also exist in DDMB binding. In Trp342 (helix X) forms a hydrogen bond with C′1 O atom and C′2-OH on the glucosyl ring; Lys18 adds one more H-bound C4-OH of galactosyl moiety; Tyr120 adds one more bifurcated H-bond to Asp19, and C6-OH is at a longer distance to Asp124. Helices are shown in ribbon. Few more sidechains including Cys27 (Helix I), Ser153 and ala 160 (helix V), and Asn244/248/251(helix VII), and Asn335 and Thr338 (helix X) are at a distance of less than 6 Å to the glucosyl moiety or the tail.

Superposition of the two structures clearly reveals that the binding residues that recognize the galactosyl moiety form a narrow pocket in MelB, named as galactoside specificity determinant pocket (**Figs. 4)**. Within the pocket, a salt-bridge-assisted H-bond network and Ch-π interactions play critical roles in the galactoside recognition. The ligand interactions are concentrated within the specificity groove, and the sugar cavity is large enough to accommodate a chain with 12 carbons. This sugar-binding site is characteristic of a narrow specificity pocket connected by a large nonspecific binding-cavity.

### Proposed cation-binding pocket

Asp55 and Asp59 on helix II have been functionally determined to be the Na^+^-binding residues to dictate the Na^+^ coordination^(22, 34, 43, 44)^. In both structures, position-59 clearly does not contact with sugar directly, but the D59C is only at 6.9-Å distance apart from C6-OH on the specific galactosyl moiety **(Fig. 5a)**. Compared to the one helical-turn apart Asp55, which is salt-bridged with Lys377 (helix XI), the position-59 is less solvent accessible (**Fig. 5b)**. The sidechain on Lys377 is only resolved to Cε position due to poor electron density. It could switch between a salt-bridge with Asp55 and a H-bond with C6-OH on the sugar. Thr121 and Gly117 on helix IV are located at one or two helical turns apart from the sugar-binding residue Asp124. The OH on Thr121 and backbone O atom of Gly117 together with the two negatively charged Asp55&59 could form a negative electrostatic surface on one side of the potential cation-binding cavity shaped by helices II, IV and XI, as well as C6-OH from the sugar. This empty site appears not completely formed; the H-bonded Thr373 and Asp124 lying on the opposite surface of this cavity could potentially contribute to both sites simultaneously when MelB at a different state, or contribute to Na^+^ binding when the sugar is absent. The obtained cooperative binding shifting to higher affinity state with the WT could be achieved through conformational changes that brings the two sites closer which allows scope for more interactions including a potential direct contact between sugar and cation. As clearly demonstrated by the experimental results, an intact cation site should be expected for this mutant. Nevertheless, the intimate physical connection between the two sites provides an essential structural basis for cooperative binding.

**Fig. 5.**
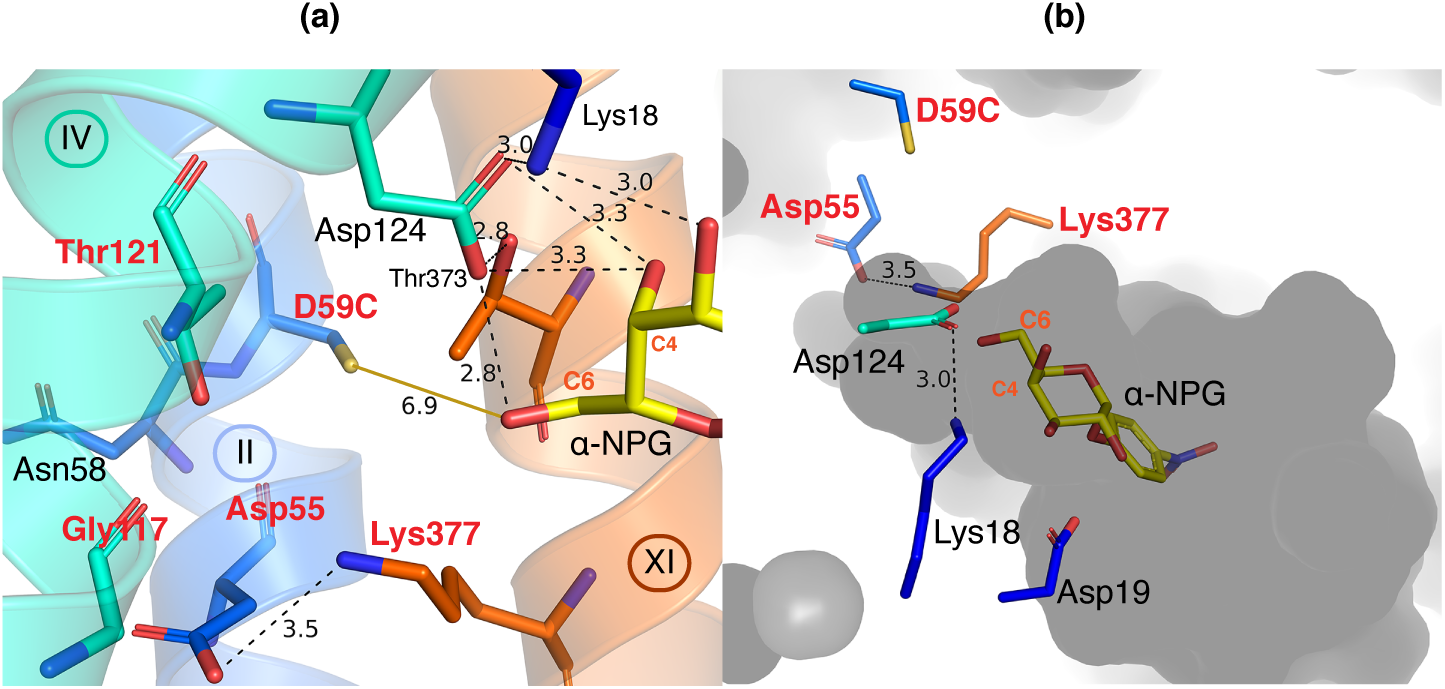
Proposed cation-binding pocket [PDB ID, 7L17]. H-bond and salt-bridge interactions are indicated by black dashed lines with distance shown in Å. **(a) Potential cation-binding site**. Positions that have been determined to be involved in MelB cation binding including Asp55 and Asp59, Gly117, Thr121, and Lys377 are labeled in red. Asp55 forms a salt-bridge with Lys377 at a distance of 3.5 Å. The C6-OH on the specific galactosyl ring is at a 6.9 Å distance to the sulfur atom on D59C, as indicated by the yellow line. Helices IV and XI carrying Tyr120, Asp124, Thr373, and Lys377 link the galactoside specificity determinant pocket to the cation pocket. **(b) A cross-section of surface representation**. Internal cavity is shown in gray. Important charged residues (4 Asp and 2 Lys) and α-NPG molecule are shown in sticks. D59C is in a less solvent-accessible position than all other sidechains.

The bioinformatics analyses show that the positions involved in the cation site are highly conserved across nearly all members analyzed, including those for different substrates, such as phospholipids with MFSD2A and MFSD2B **(SI Fig. S5)**. On the other hand, the galactoside-binding residues are only conserved within MelB orthologues from varied bacterial strains.

## Discussion

Results presented here or in the literatures^(7, 44)^ show that D59C MelB mutant selectively abolishes the cation binding and all three modes of cation-coupled melibiose transport, but remains capable of galactoside binding and melibiose translocation^(7, 34, 44)^. This offered an opportunity to simplify the coupled symport to an uncoupled uniport. Crystallization of the WT MelB with bound sugar is quite challenging, probably due to the greater dynamic nature and more conformational states of being a symporter. With this uniport mutant carrying only sugar binding and translocation properties, the protein is more stable and crystallization trials are much more reproducible.

Overlay of the two structures with bound α-NPG or DDMB clearly reveals the galactoside specificity determinant pocket is responsible for the galactoside recognition and binding in MelB (**Figs. 2-4**). The helices II (Lys18, Asp19, Ile22, and Tyr26, particularly Asp19 and Lys18) and VI (Tyr120, Asp124, and Trp128, particularly Asp124) make major contributions to the sugar binding, and helices V (Arg149), X (Trp342), and XI (Thr373 and Val376) also contributes to the binding affinity. Mutations on most of these positions largely reduce the binding or transport in MelB_St_^(7)^ or MelB_Ec_, and some affected affinity to both sugar and cation^(34, 47-50)^. The structure was also strongly supported by the Fourier-transform infrared spectroscopy (FTIR) studies in MelB_Ec_, where the mutant on positions-19 or -124 lost sugar-specific FTIR signals and D124C also largely reduced the Na^+^-specific signals^(34)^.

There are only few disaccharide transporters with structure solved^(51-53)^. Similar to the sugar-binding site in LacY^(51)^ and maltose-binding protein MalE in the maltose transporter MalEFGK_2_ ^(54)^, charged residues dictate the sugar binding, but it is not the case for the low-affinity maltose site within the membrane-domain in MalF^(53)^. Interestingly, an “acceptor-donor-acceptor” pattern for H-bonding with sugar is also found in MalE^(54)^. Aromatic residues presented in these sugar-binding sites form CH/π-interactions between aromatic and the galactosyl rings, or aromatic stacking interactions with phenyl ring, which shapes the binding pocket and increases the binding strength^(55, 56)^. Different from LacY, the MelB sugar-binding site displays a comprehensive salt-bridge-assisted H-bonding network; the sugar arrangement in both sites is also different. While both permeases transport galactosides, the sugar OH position connecting to its coupling cation site also differs. The C3 and C2-OH groups link with LacY H^+^ site^(51, 52)^, while the C6-OH is in close proximity to MelB cation site **(Fig. 5)**.

The two structures with bound α-NPG and DDMB can explain well why MelB can recognize a large variety of galactosides from mono- to tri-saccharides, as well as other glycoside analogues (methyl-thiogalactosides, methyl-galactosides, NPG), dansyl-galactoside with 2 to 6 carbons between the galactosidic and the dansyl moieties^(24, 26, 33)^. A galactosyl moiety that is conserved among all sugars for recognition by the selection pocket, and the chemical variation on the rest non-galactosyl moiety can be accommodated nonspecifically by a large cavity. The nonspecific interactions largely increase the binding affinity. This structural insights into the galactoside binding might suggest a potential drug delivery strategy based on glycosides being transported by a sugar active transport.

Determination of the sugar-binding site in MelB allows to recognize the nearby cation specificity determinant pocket (**Fig. 5**), which confirms the previous conclusion based on a large body of biochemical/biophysical analyses including mutagenesis and second-site suppressors, all favoring an overlapping sugar- and cation-binding sites^(34, 47, 48, 50, 57, 58)^. While no Na^+^ bound with MelB, several positions including 55, 59, 117 and 121 have been determined essential or important for the binding of Na^+^ and Li^+^ and coupled transported^(7, 22, 34-36, 43, 44, 59, 60)^. Asn58 at the neighboring position of 59 play important role in the cation selectivity because Ala58 in *Klebsiella pneumonia* MelB selectively removes the Na^+^ recognition and retains the H^+^- and Li^+^- coupled transport^(61, 62)^. It is also noteworthy that Arg replacement on Gly117 decreases affinity for Na^+^ and Li^+^ by about 10-fold, and failed in melibiose active transport and efflux, but catalyzes melibiose exchange coupled to Na^+^ or Li^+^ at a normal rate^(36)^. Cys replacement on Thr121 selectively inhibit Na^+^-coupled melibiose transport with less effect on Li^+^-coupled melibiose transport^(7)^. On the homologue site in human MFSD2A, the substitution of Met for Thr results in a lethal mutation^(63)^. For another human homologue MFSD13, identical fragments that host the four positions (^55^<u>D</u>AI<u>ND</u>^59^ and ^117^<u>G</u>MTY<u>T</u>IM<u>D</u>^124^) are observed **(SI Fig. S6b)**. It is likely that those human homologues are cation-coupled symporters as demonstrated in MFSD2A^(3)^.

Functionally, all three cations (H^+^, Na^+^, or Li^+^) compete for the same site with a stoichiometry of unity. While a metal site could be conceptualized based on the structure, where is the site for H^+^? Competitive Na^+^ binding at varied pH revealed a p*K*a of 6.25 or 6.59 for MelB_St_ in the absence or presence of melibiose^(22)^, and p*K*a of 6.3 for MelB_Ec_ ^(64)^. The p*K*a for Asp55 is expected to be at a normal range of around 3.6 and the salt-bridged Asp55 with Lys377 are solvent accessible. Asp59 lies in a less-solvent exposed position and an elevated p*K*a for this carboxyl group is expected. More direct evidence could be that D59C MelB_Ec_ lacks transient alkalization of medium during melibiose gradient-driven melibiose transport^(44)^. Thus, Asp59 is the H^+^-binding residue in MelB, and the D59C mutant lacks affinity for all three cations.

The position 59 regulates MelB between symport and uniport for melibiose transport; a transport kinetic model for both can be simplified as an 8-step or 6-step cycle, respectively **(Fig. 6)**. For the uniport process, melibiose is recognized [i] by the apo mutant, which induces alternating-access mechanism and shifts MelB conformation into an occluded intermediate [ii] and opens to the cytoplasm [iii]. Bound melibiose is released into the cytoplasm [iv], and MelB closes the cytoplasmic opening [v] and resets to the outward-open state [vi] followed by a new cycle. Since the *K*_d_ value for D59C mutant is as low as 6 mM, it can only work when environmental melibiose at high concentrations, such as 30 mM used in the MacConkey agar or in melibiose exchange experiment^(7)^. Under active transport assay, melibiose was presented at 0.4 mM, so no uptake is detected **(Fig. 1)**. At step [iv], melibiose release or rebound is dependent on the intracellular melibiose concentration. In the fermentation assay, the incoming melibiose is immediately hydrolyzed by a high-turnover number enzyme α-galactosidase, so no melibiose concentration can be built up. Melibiose cytoplasmic release is favored over rebound. For the symporter, two more steps are added including cation binding and release **(Fig. 6b)**, which allow MelB to work at environmental low melibiose conditions by increasing the melibiose binding through a positive cooperativity with a coupled cation^(22)^. In the presence of electrochemical ion gradience of H^+^, Na^+^, or Li^+^, the cytoplasmic release of bound cation at step [6] is favored, which will facilitate the cytoplasmic closure and prevents the melibiose rebound, so collectively, the transport *K*_m_ can be largely improved^(27, 35, 65)^.

**Fig. 6.**
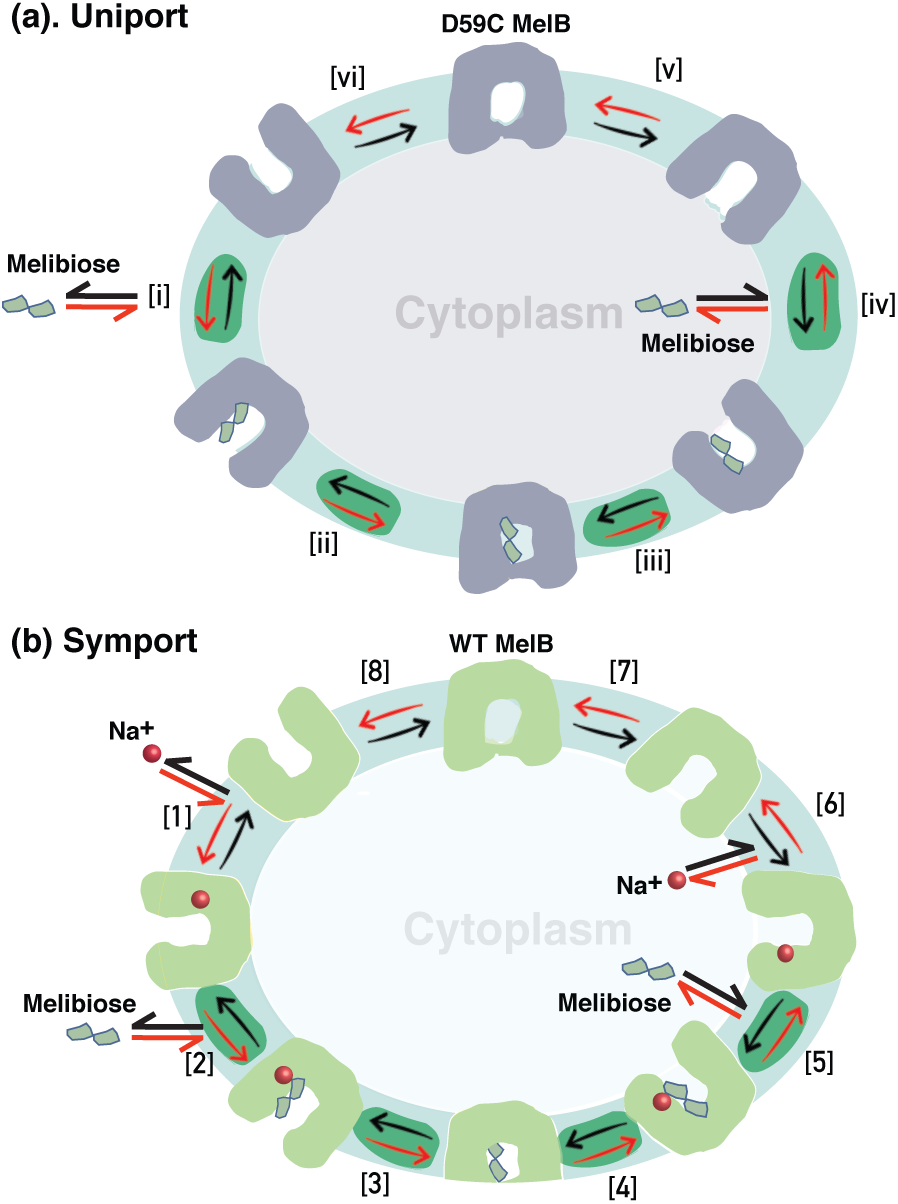
Models for the symport and uniport of melibiose transport. The symbols for the WT and D9C MelB proteins, Na^+^, and melibiose are indicated. **(a) Uniport catalyzed by the D59C MelB mutant**. Melibiose inward-directly transport begins at step i and proceeds via the red arrows around the circle of 6 steps, with one melibiose across the membrane per circle. Melibiose efflux transport begins at step iv and proceeds via the black arrows around the circle, with one melibiose across the membrane per circle. Melibiose exchange begins at step iv, and takes one intracellular melibiose out in exchange with extracellular melibiose molecule, only involved in step iv – i, a 4-step reaction as highlighted in green color. **(b) Symport catalyzed by WT MelB**. Melibiose active transport begins at step 1 and proceeds via the red arrows around the circle, with one melibiose and one cation inwardly across the membrane per circle. Melibiose efflux transport begins at step 6 and proceeds via the black arrows around the circle, with one melibiose and one cation outwardly across the membrane per circle. Melibiose exchange begins at step 6, and also takes 4 steps involved step 5 - 2 as highlighted in green color.

In summary, this structural study reveals that the specificity determinants in MelB for galactosides are within a narrow groove that connects to the cation-binding pocket. The intimate connection between the two specificity determinant pockets lays the structural basis for the obligatory coupling of this family of transporters. The sugar recognition in MelB characterized by a specificity groove along with the large nonspecific cavity to accommodate structurally diverse non-galactosyl moieties offers a potential for future exploration of active transporters for drug design and delivery.

## Materials and Methods

### Materials and Reagents

[1-^3^H]Melibiose was custom-synthesized by PerkinElmer (Boston, MA). Other chemicals and reagents used in this study were of analytical grade and purchased from standard commercial sources except 2′-(*N*-dansyl)aminoalkyl-1-thio-β-D-galactopyranoside (D^2^G) that was kindly provided by Drs. Gerard Leblanc and H. Ronald Kaback. The detergents undecyl-β-D-maltopyranoside (UDM), dodecyl-β-D-maltopyranoside (DDM), and dodecyl-β-D-melibioside (DDMB), and *E. coli* polar lipids (Extract Polar, Avanti,100600) were purchased from Anatrace. Melibiose and 4-nitrophenyl-α-D-galactopyranoside (α-NPG) were purchased from Sigma-Aldrich. The SelenoMet Media were purchased from Molecular Dimensions Limited. Crystallization reagents and materials were purchased from Hampton Research.

### Plasmids and cell culture for transport assays

The overexpression of MelB_St_ was carried out in the *E. coli* DW2 strain (*mel*A^+^, *mel*B, and *lac*ZY) from a constitutive expression plasmid pK95ΔAH/MelB_St_/CHis_10_ ^(7, 26, 31)^. The D59C mutant with Cys at the position 59 was constructed previously^(7)^. *E. coli* DW2 cells containing a given plasmid were grown in Luria-Bertani (LB) broth (5 g yeast extract and 10 g tryptone per liter with 171 mM NaCl) with 100 mg/liter of ampicillin in a 37 °C shaker. The overnight cultures were diluted by 5% with LB broth supplemented with 0.5% glycerol and 100 mg/liter of ampicillin, and constitutive overexpression was obtained by shaking at 30 °C for another 5 h.

### Melibiose fermentation

The DW2 cells were transformed with a given plasmid, plated on MacConkey agar plate supplemented with melibiose at 30 mM (the sole carbohydrate source) and 100 mg/liter ampicillin, and incubated at 37 °C. After 18 h, the plates were viewed and photographed immediately.

### [1-^3^H]Melibiose transport assay

*E. coli* DW2 cells expressing MelB_St_ were washed with 100 mM KP_i_, pH 7.5 to remove Na^+^ contamination as described previously^(26)^. The cell pellets were resuspended with the assay buffer (100 mM KP_i_, pH 7.5, 10 mM MgSO_4_) and adjusted to *A*_420_ of 10 (∼0.7 mg protein/ml). Flux assay with a fast filtration was carried out to analyze the transport time course, and intracellular melibiose was determined as described previously.

### Large-scale protein production

LB broth supplemented with 50 mM KPi (pH 7.0), 45 mM (NH_4_)SO_4_, 0.5% glycerol, and 100 mg/L ampicillin was used for fermentation, and protocols for membrane preparation and MelB_St_ purification by cobalt-affinity chromatography after extracted in detergent UDM have been described previously^(7)^. MelB_St_ protein solutions were dialyzed overnight against a buffer (consisting of 20 mM Tris-HCl, pH 7.5, 100 mM NaCl, 0.035% UDM, and 10% glycerol), concentrated with Vivaspin column at 50 kDa cutoff to approximately 40 mg/ml, and subjected to ultracentrifugation at 384,492□*g* for 45□min at 4□°C (Beckman Coulter Optima MAX, TLA-100 rotor), stored at -80 °C after flash-frozen with liquid nitrogen.

### Seleno-methionine incorporation

The seleno-methionine derivative of MelB_St_ D59C mutant (SeMet D59C MelB_St_) was obtained by incorporating Sel-Met during the protein expression in the *E. coli* strain DW2 in the SelenoMet Medium. When supplemented Met at a ratio of 10:1 for Sel-Met vs. Met to the medium, the SeMet D59C MelB_St_ sample failed to produce crystals, so partial labeling at a ratio of 3:1 (66.7% sel-Met) was applied.

### Protein concentration assay

The Micro BCA Protein Assay (Pierce Biotechnology, Inc.) was used for the protein concentration assay. Protein concentration for crystallization trails were estimated by measuring the UV absorption at 280 nm.

### Circular dichroism (CD) spectroscopy

MelB_St_ at 10 μM in 10 mM NaPi, pH 7.5, 100 mM NaCl, 10% glycerol, and 0.035% UDM was analyzed with Jasco J-815 spectrometer equipped with a peltier MPTC-490S temperature-controlled cell holder unit. The CD spectra for a wavelength range of 200-260 nm were carried as described previously^(46)^. Melting temperature (*T*_m_) determination was carried out at temperatures between 25 - 80 °C. Ellipticity at 210 nm were recorded at a 1 °C interval with the temperature ramp rate at 1 °C per minute, and plotted against temperature. The *T*_m_ values were defined as the temperature leading to the half maximal decrease in the helical content, which was determined by fitting the data using the Jasco Thermal Denaturation Multi Analysis Module.

### Isothermal titration calorimetry (ITC) measurements

All ITC ligand-binding assays were performed with the TA Instruments (Nano-ITC device). In a typical experiment, the titrand (MelB_St_) in the ITC sample cell was titrated with the specified titrant (placed in the Syringe) by an incremental injection of 2-µL aliquots at an interval of 300 sec at a constant stirring rate of 250 rpm (nano-ITC). All samples were degassed using a TA Instruments Degassing Station (model 6326) for 15 min prior to the titration. Heat changes were collected at 25 °C, and data processing was performed with the NanoAnalyze (version 3.6.0 software) provided with the ITC equipment. The normalized heat changes were subtracted from the heat of dilution elicited by last few injections, where no further binding occurred, and the corrected heat changes were plotted against the molar ratio of titrant versus titrand. The values for association constant (*K*_a_) value were determined by fitting the data with a one-site independent-binding model. At most cases, the binding stoichiometry (*N*) number was fixed to 1 since it is a known parameter, which can restrain the data fitting and achieve more accurate results^(66)^. The dissociation constant *K*_d_ = 1/*K*_a_.

### Trp→dansyl fluorescence resonance energy transfer (FRET) and IC_50_ determination

Steady-state fluorescence measurements were performed with an AMINCO-Bowman series 2 spectrometer with purified MelB_St_ at 1 μM in 20 mM Tris-HCl, pH 7.5, 100 mM CholCl, 10% glycerol, 0.03% UDM. Using an excitation wavelength at 290 nm, the emission intensity was recorded at 490 nm. On a time trace, D^2^G, NaCl or LiCl, then DDMB or α-NPG were sequentially added into the protein solution at a 1 min-interval. For the determination of IC_50_, after the addition of 10 μM D^2^G (the *K*_d_ for the WT), DDMB, DDM, UDM, or α-NPG was consecutively added until no change in fluorescence intensity occurred. An identical volume of water was used for the control. The decrease in intensity after each addition of DDMB or α-NPG was corrected by the dilution effect and plotted as a function of the accumulated DDMB or α-NPG concentrations. The 50% inhibitory concentration (IC_50_) was determined by fitting a hyperbolic function to the data (OriginPro 2020b).

### Crystallization, native diffraction data collection and processing

Crystallization trials were carried out by the hanging-drop vapor-diffusion method at 23□°C by mixing 2□μl of the pre-treated protein samples containing α-NPG or DDMB with 2□μl reservoir. To prepare the α-NPG-containing D59C MelB_St_, the protein samples in the dialysis buffer composed of 20 mM Tris-HCl, pH 7.5, 100 mM NaCl, 0.035% UDM, and 10% glycerol was diluted to 10 mg/ml and supplemented with phospholipids at a concentration of 3.6□mM (*E. coli* Extract Polar, Avanti, 100600) from a 20-mM stock dissolved with a dialysis buffer containing 0.01% DDM instead of 0.035% UDM, as well as 6 mM α-NPG. To prepare DDMB-containing D59C MelB_St_, the same treatment was carried out except that the protein samples were supplemented with 0.015% DDMB (1 x CMC) and 10% PEG3350. All samples were incubated for 15 min prior to the crystallization trials, and both crystals of D59C MelB_St_ containing α-NPG or DDMB were grown using a reservoir consisting of 50 mM BaCl_2_, 50 mM CaCl_2_, 100 mM Tris-HCl, pH 8.5, and 29-32% PEG 400. The crystals can be obtained from a wide range of reservoir conditions with no notable change on the structure; such as, 50 mM BaCl_2_ can be replaced by 100 mM NaCl, or 100 mM Tris-HCl, pH 8.5 can be replaced by 100 mM MES, pH 6.5. Crystals appeared in 3-4 days, were frozen with liquid nitrogen in 2-3 weeks, and tested for X-ray diffraction at the Lawrence Berkeley National Laboratory ALS BL 5.0.1 or 5.0.2 via remote data collection method. The complete diffraction datasets for α-NPG- or DDMB-bound native crystals were collected at 100□K from a single cryo-cooled crystal at a wavelength of 0.97949 Å on ALS BL 5.0.2 or 0.97741□Å on ALS BL 5.0.1, respectively, with a Pilatus3 6M 25 Hz detector. ALS auto-processing XDS output files were used for structure solution. The statistics in data collection is described in **Table 3**.

**Table 3.** Crystallographic Data collection, phase, and refinement statistics

| **Data collection** | SelMet D59C MelB<sub>St</sub><br>with DDMB | D59C MelB<sub>St</sub> with α-NPG<br>[PDB ID, 7L17] | D59C MelB<sub>St</sub> with DDMB<br>[PDB ID, 7L16] |
| --- | --- | --- | --- |
| Space group | P 31 2 1 | P 31 2 1 | P 31 2 1 |
| Cell dimensions |  |  |  |
| *a*, *b*, *c* (Å) | 126.4 126.4 103.7 | 126.364 126.364 104.334 | 127.372 127.372 103.905 |
| $\alpha$, $\beta$, $\gamma$ (°) | 90 90 120 | 90 90 120 | 90 90 120 |
| Resolution (Å) | 38.4 - 4.0<sup>#</sup> | 29.89 - 3.05* | 48.72 - 3.15* |
| *R*<sub>sym</sub> or *R*<sub>merge</sub> | 0.185 (1.213) | 0.111 (1.404) | 0.138 (1.463) |
| *I* / $\sigma I$ | 77.4 | 20.5 (2.4) | 18.2 (2.6) |
| Completeness (%) | 99.41 | 99.48 (99.95) | 98.66 (100.00) |
| Redundancy | 64.4 (50.51) | 21.6 (22.5) |  |
| **Refinement** |  |  |  |
| Resolution (Å) |  | 29.35 - 3.05 (3.159 - 3.05) | 29.35 - 3.15 (3.263 - 3.15) |
| No. reflections |  | 18614 (1862) | 16973 (1684) |
| *R*<sub>work</sub> / *R*<sub>free</sub> |  | 0.254 / (0.287) | 0.273 / 0.297 |
| No. atoms |  | 3582 | 3596 |
| Protein |  | 3561 | 3561 |
| Ligand/ion |  | 21 | 35 |
| Water |  | 0 | 0 |
| *B*-factors |  | 101.91 | 103.07 |
| Protein |  | 101.94 | 102.92 |
| Ligand/ion |  | 96.56 | 117.89 |
| R.m.s. deviations |  |  |  |
| Bond lengths (Å) |  | 0.003 | 0.002 |
| Bond angles (°) |  | 0.615 | 0.520 |

<sup>#</sup> Merged from 13 datasets; \*A single crystal was used for the structure. Values in parentheses are for highest-resolution shell.

### Anomalous diffraction data collection, processing, and experimental phasing

The crystals from SeMet D59C MelB_St_ mutant were obtained from 50 mM BaCl_2_ or 100 mM NaCl, 50 mM CaCl_2_, 100 mM Tris-HCl, pH 8.5, or 100 mM MES, pH 6.5. Weak anomalous signals presented in each individual dataset, and multiple datasets at varied wavelength were collected and processed either manually by HKL2000^(67)^ or DIALS in CCP4i2^(68)^, or using the ALS auto-processing DIALS method. Phenix Scale and Merge Data and Phenix Xtriage programs (v.1.18.2-3874 or de-3936)^(69)^ were used for selecting and merging dataset. A merged dataset, which was generated from 13 datasets collected from 6 crystals with a resolution cutoff between 40 - 4.0 Å, exhibits anomalous signal with CC_ano half_ >0.94 at 8.0 Å and 0.04 at 4 Å, as analyzed by Phenix Anomalous Signal program. Among those data sets, most were from a suboptimal wavelength for Se atom (0.977408 Å) collected on BL5.0.1, and one from a wavelength of 0.97919 Å on BL5.0.2.

Searching for selenium atoms and phasing at a several resolutions limited to 6.0 to 6.5 Å (CC_ano_ half >0.44) were performed using ccp4i2 (7.0) Crank2 Phasing and Building program^(70)^ and Phenix Hybrid Substructure Search program^(71, 72)^. A total of 18 selenium sites were identified, and used for phasing and model auto-building by SAD or MR-SAD using a model derived from the pdb id 4M64 Crank2 Phasing and Building programs. The model building was performed in Coot 0.8 or 0.9^(73)^ and refined against the combined anomalous dataset at 4.0 Å by Phenix Refinement^(74)^.

### Structure determination of native D59C MelB_St_ and α-NPG or DDMB fitting

The structure determination for the native data sets was performed by molecular replacement used the refined SeMet model as the search model in the Phenix Phaser-MR^(71)^, and followed by rounds of manual building and refinement. Overlay of the 18 sites of selenium atoms is shown in the **SI Fig. S4**. Ligand fit using α-NPG 3D structure with a ligand code 9PG from pdb id 4zyr ^(51)^ was performed with Phenix LigandFit program^(75)^ for the well-refined models. The 3-D structure of DDMB is not available, so a 2D-structure was drawn by Chemdraw; Phenix eBLOW^(76)^and Readyset were used to generate and optimize the ligand restraints for refinement. The manually aligned DDMB structure on α-NPG was used for LigandFit. All ligand-bound structures were modeled from positions 2 to 454 without a gap. After rounds of refinement, the structures with α-NPG or DDMB were refined to a resolution of 3.05 Å (PDB access ID, 7L17] or 3.15 Å [PDB access ID, 7L16], respectively. Several crystal structures of D59C mutant bound with α-NPG or DDMB were refined to resolutions beyond 3.3 Å, and all structures are virtually identical. The first met was not present as also shown by N-terminal sequencing analyses with MelB_St_ (Alphalyse Inc. CA). The statistics of refinement for the final models are summarized in **Table 3**. For α-NPG- and DDMB-bound structures: Ramachandran favored of 98.67% and 95.57%, Ramachandran outliers of 0%, clashscores of 2.48 and 2.19, and overall scores of 1.05 and 1.30, respectively, as judged by MolProbity in Phenix. Pymol (2.3.5) was used to generate all graphs.

### Statistics and reproducibility

All experiments were performed 2-4 times. The average values were presented in the table with standard errors.

## Acknowledgements

The authors thank Drs. R. Bryan Sutton, Luis Cuello, Abdul Ethayathulla, and Kei Nanatani for discussions and helps. The strain DW2 and the expression vector were from Dr. Gerard Leblanc and reagent D^2^G was from Drs. Gerard Leblanc and Ronald Kaback. This study is supported by the National Institutes of Health grants R01GM122759 and R21NS105863 to L.G.

## Author contribution

L.G. designed this study and performed protein crystallization, X-ray diffraction data collection and processing, structure determination, and interpretation. P.H. performed all protein purification, functional characterizations and related data processing and figure preparations. L.G. wrote the manuscript with help of P.H.

## Competing interests

The authors declare no competing financial interests.

## Additional Information

### Accession codes

The protein coordinates and structure factors have been deposited in the Protein Data Bank with the accession numbers 7L16 and 7L17 for the D59C MelB_St_ bound with DDMB or α-NPG, respectively.

## Supporting information figure legend

**SI Fig. S1. ITC measurements**. ITC measurements were performed at 25 °C as described in Methods. **(a) Binding thermography**. For all tests, ligands and proteins are buffer-matched. For melibiose binding measurements, a solution of 80 mM or 10 mM in the absence or presence of NaCl at 100 mM, was injected into the ITC sample cell containing the WT MelB_St_ at 80 μM without or with 100 mM NaCl, respectively. With the D59C mutant, 40 melibiose was used. For α-NPG binding measurements, 0.5 mM or 0.4 mM solution was used to titrate the WT or D59C mutant at 50 μM in the absence or presence or 100 mM NaCl, respectively. For the Na^+^ or Li^+^ binding measurements, 5 mM or 2 mM solution was used to titrate WT MelB_St_ at 80 μM without or with 50 mM melibiose, respectively; 40 mM solutions was used to titrate the D59C mutant. **(b) binding isotherm**. Curve fit was carried out with a one-site independent-binding model included in the NanoAnalyze software (version 3.6.0). Results are presented in **Table 1**.

**SI Fig. S2. Determination of DDMB binding affinity to MelB**_**St**_. **(a) Titration with DDMB or** α**-NPG**. MelB_St_ in 20 mM Tris-HCl, pH 7.5, 100 mM CholCl, 10% glycerol, 0.03% UDM without (1) or with 100 mM NaCl (2) were added with D^2^G at 10 μM at 1-min time point as indicated by triangle symbols on the Trp→dansyl FRET time trace. Starting at the 2-min of time point, DDMB or α-NPG solution was supplemented consecutively at a 1-min interval into MelB_St_ solution till no change in fluorescence intensity reached as indicated in star symbols. DDM and UDM were used as negative controls. The accumulated function concentration for both is less at 300 μM, which is close the DDMB CMC value. At 2-min time point, additions of galactosides were colored in pink, α-NPG in yellow, UDM in blue, DDM in light blue, water in gray traces, respectively. **(b) IC50 determination**. The intensity decrease at each titration were corrected by the dilution effect, and plotted as a function of the accumulated DDMB or α-NPG concentration. The 50% inhibitory concentration (IC_50_) was determined by fitting the data to a hyperbolic function (OriginPro 2020b). Results are presented in **Table 2**.

**SI Fig. S3. CD spectra and *T***_**m**_ **determination**. MelB_St_ at 10 μM in 10 mM NaPi, pH 7.5, 100 mM NaCl, 10% glycerol, and 0.035% UDM was used for the CD analysis as described in Methods. **(a) CD spectra**. The CD was recorded between 200 – 260 nm for both the WT and D59C mutant MelB_St_ in the absence of sugar. **(b) Thermo-denaturation**. The tests were carried out between 25 to 80 °C in the absence of sugar. The *T*_m_ values was determined using the Jasco Thermal Denaturation Multi Analysis Module.

**SI Fig. S4. Stereo view of electron density maps of D59C MelB**_**St**_ **bound with** α**-NPG. (a)** Cross-eye stereo view. The structure for D59C MelB_St_ bound with α-NPG [PDB ID, 7L17] is shown in ribbon representation and overlayed with 2Fo-Fc electron density map contoured at 1.5 σ. Helices V and VIII are placed in the front. α-NPG is shown in sticks. **(b)** Cross-eye stereo view of selenium atoms. A total of 18 selenium atoms identified based on anomalous signal of selMet D59C MelB_St_ bound with DDMB were overlayed with refined structure of D59C MelB_St_ structure bound with α-NPG. The protein is shown in cartoon representation; 18 Met sidechains are shown in stick and ball with sulfur atom colored in yellow and carbon atoms colored matching with individual backbones. Selenium atoms are colored in black. Except for M316 and M423, all other overlayed well with selenium atoms. The bound α-NPG is shown in sticks.

**SI Fig. S5. Helical packing and overall architecture of MelB**_**St**_. The α-NPG-bound D59C MelB_St_ structure [PDB ID, 7L17] is colored in rainbow spectrum. The bound α-NPG in each panel is colored in yellow. Upper row, helical packing in cartoon representation; bottom row, surface representation. **(a-b) Side view with helices V and VIII in front or II and XI in front, respectively. (c) Viewed from cytoplasmic side. (d) Viewed from periplasmic side**.

**SI Fig. S6. Cation and sugar sites. (a) An overall view of co-substrate binding sites**. The specific determinant pocket is closely connected with the proposed cation site as labeled in red. Helices are shown in a cartoon representation and labeled in Roman numerals; α-NPG is colored in yellow with oxygen in red and nitrogen in blue. **(b) Alignment of selected MelB homologues**. MelB_St_ is aligned with other 13 bacterial orthologues that are melibiose transporters or function unknown transporters, or with three selected human transporters, respectively. Protein sequence access identifications are presented. Selected regions containing the positions involved in sugar binding and cation site are isolated and presented. The transmembrane locations are indicated according the MelB_St_ structure. Yellow color high-lights the identical positions that in MelB is in the galactosyl moiety-binding site (18, 19, 22, 26,124, 128,129, 342, 376). Except for Ile22, Tyr26, and Val376 with hydrophobic contacts, all are engaged in one to three H-bonding interaction(s) with galactosyl moiety. Pale yellow highlights the identical positions involved in the H-bonding network with no direct contact with galactoside in MelB. Color in cyan highlights the identical positions involved in cation binding in MelB. The sequence for MELB_ECO57 was renumbered by adding MSIS-at the N-terminus. It is noteworthy that “MSIS-” sequence for most MelB was missed at earlier time, which might cause a confusion; such as, Asp55 and Asp59 were named as Asp51 and Asp55, respectively.

---

**Extended Data Figure 1.**
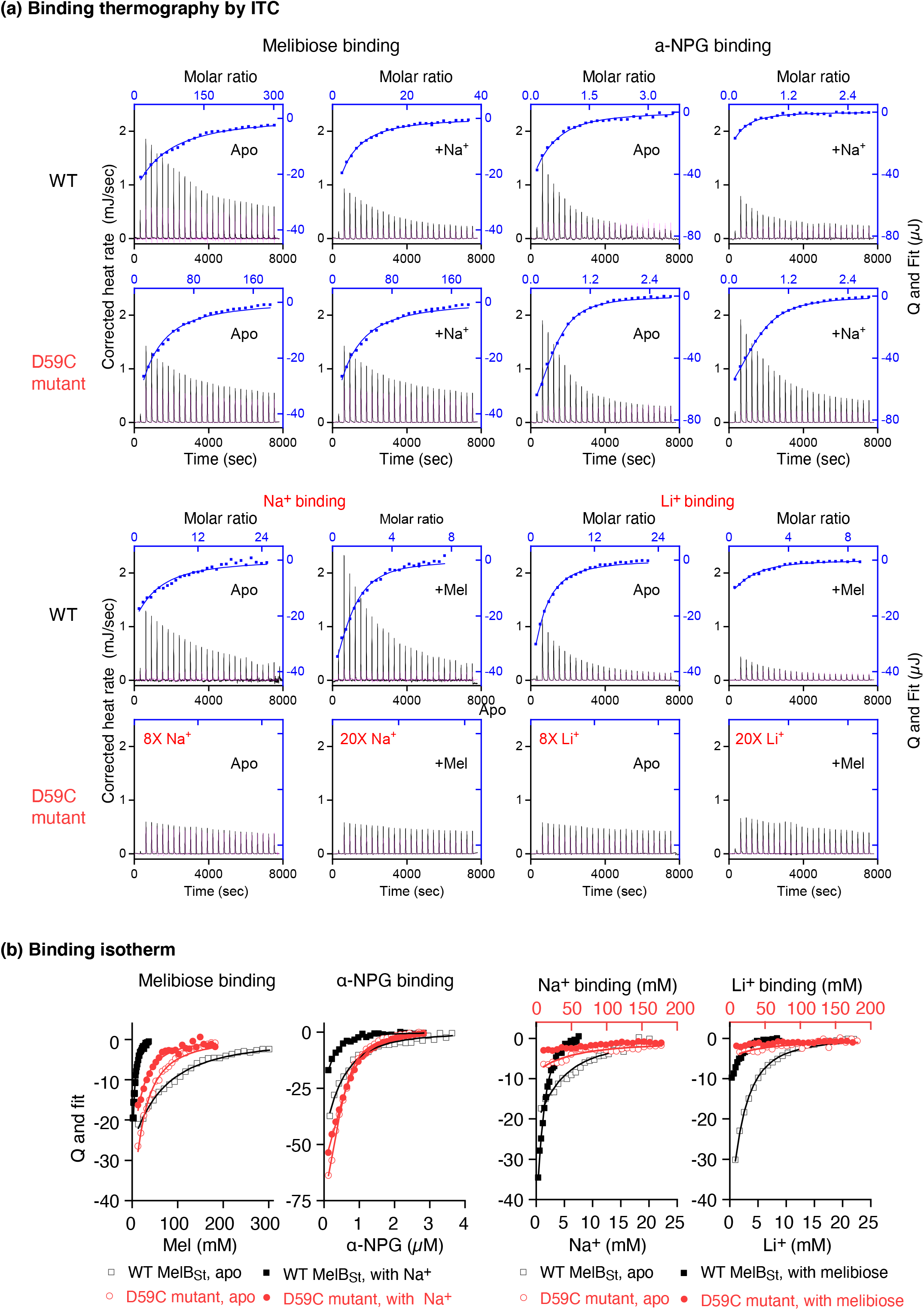

## Notes

### Competing Interest Statement

The authors have declared no competing interest.

