## Supplementary figures and images for "Molecular recognition of sugar binding in a melibiose transporter MelB by X-ray crystallography"

### SI Fig. S1

#### Supporting information Figure S1

##### (a) Binding thermography by ITC

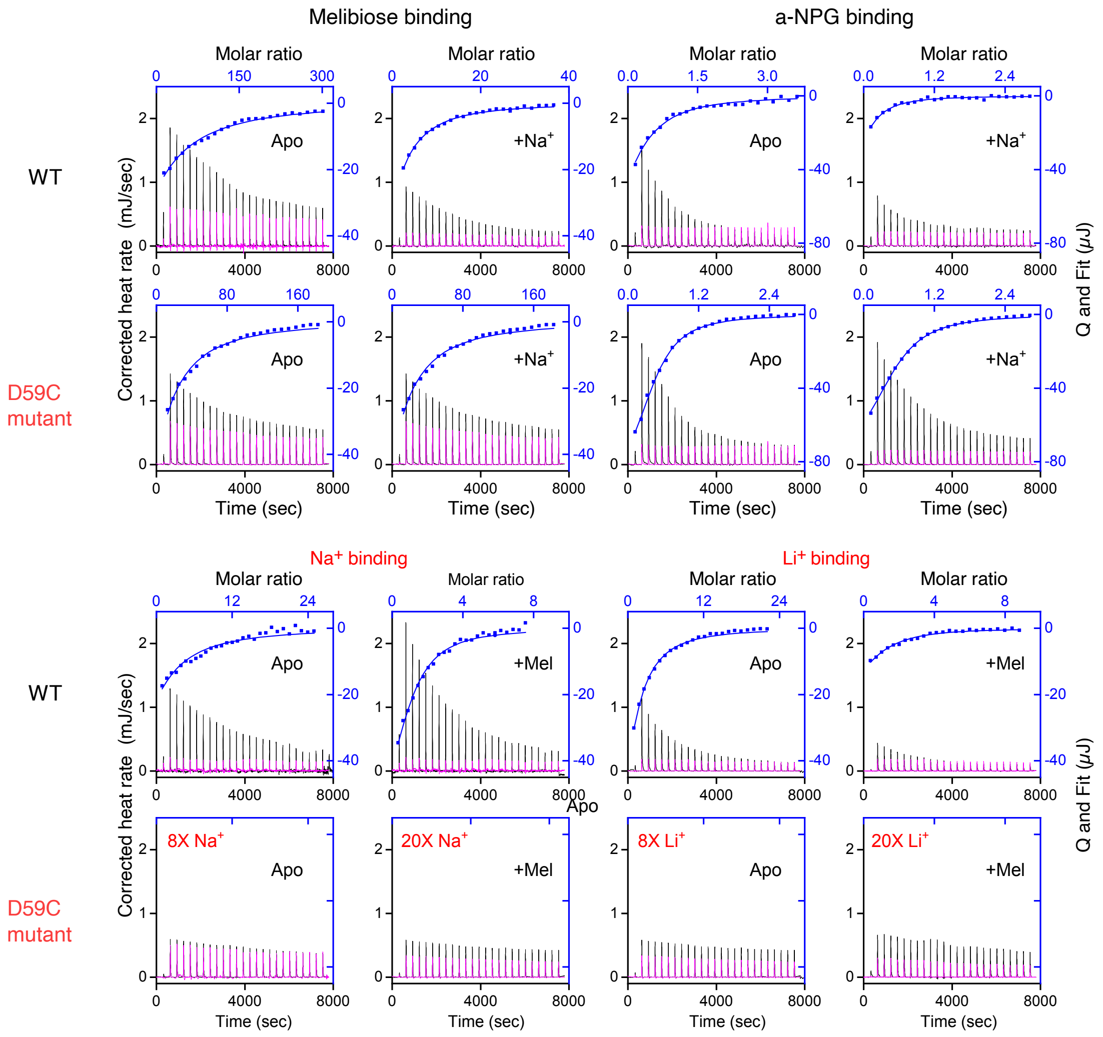


##### (b) Binding isotherm

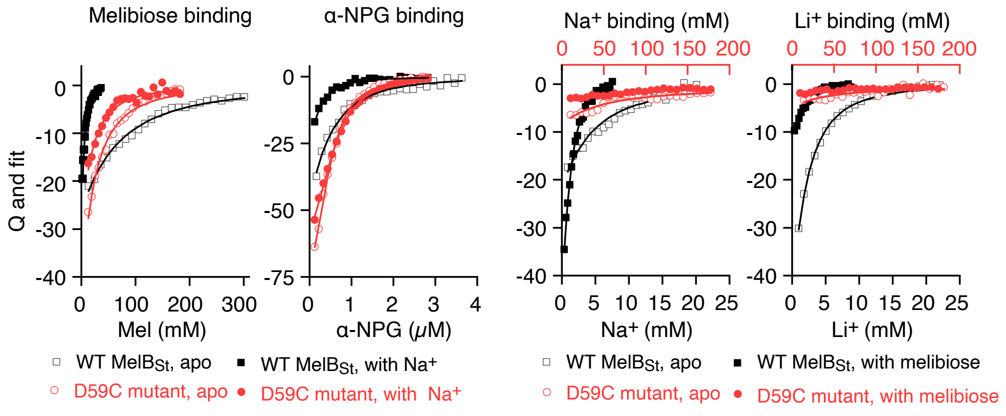

### SI Fig. S2

**Supporting Information Fig. S2**

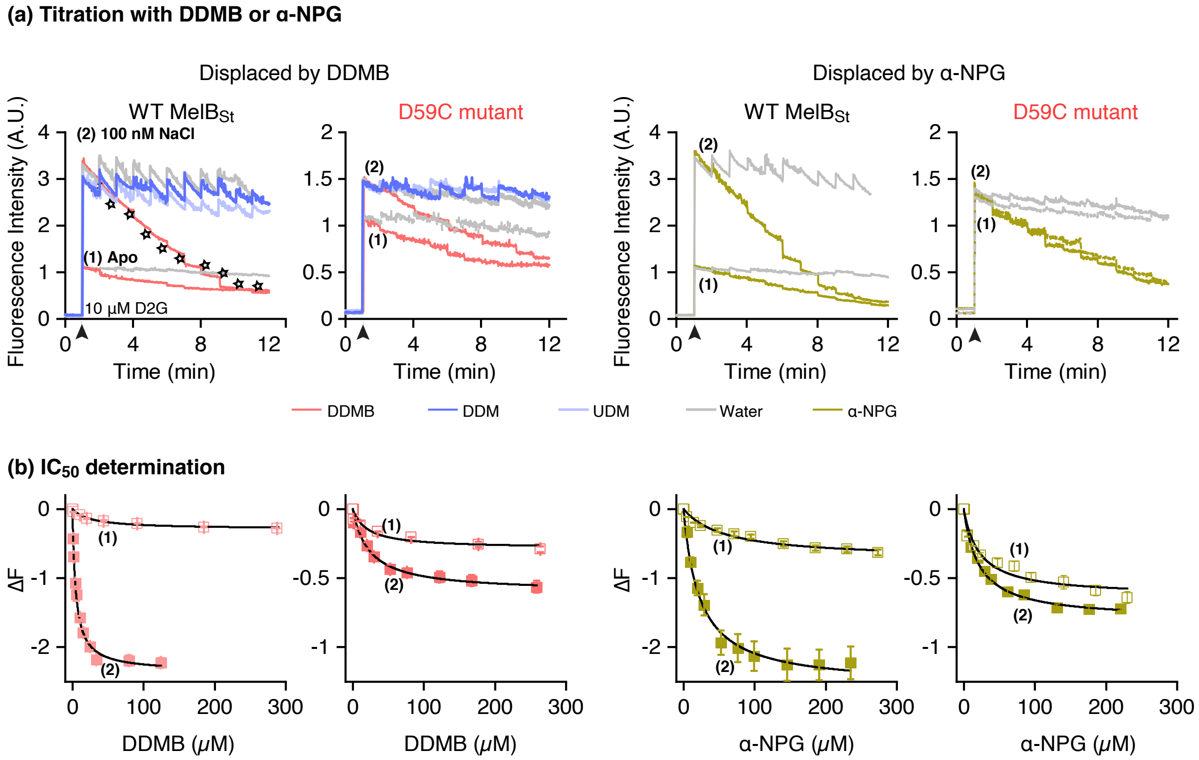

### SI Fig. S3

**Supporting Information Fig. S3**

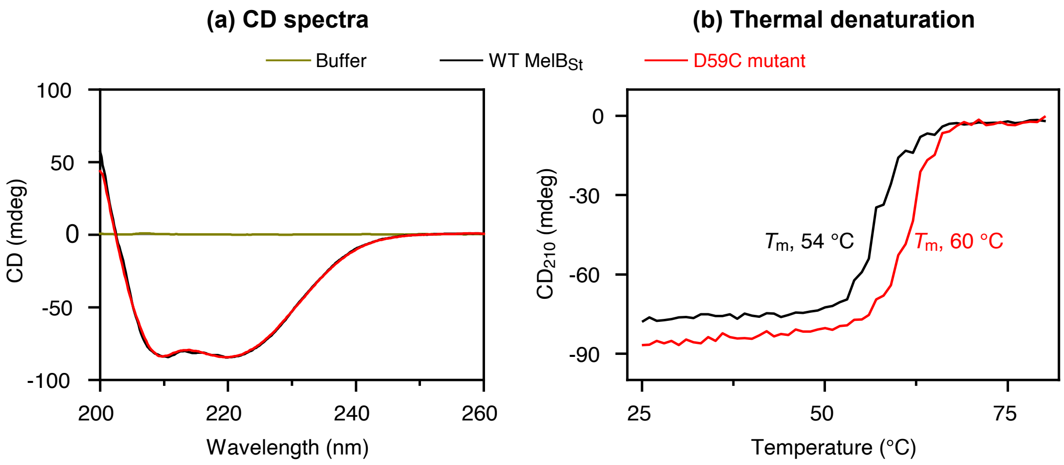

### SI Fig. S5

**Supporting Information Fig. S5**

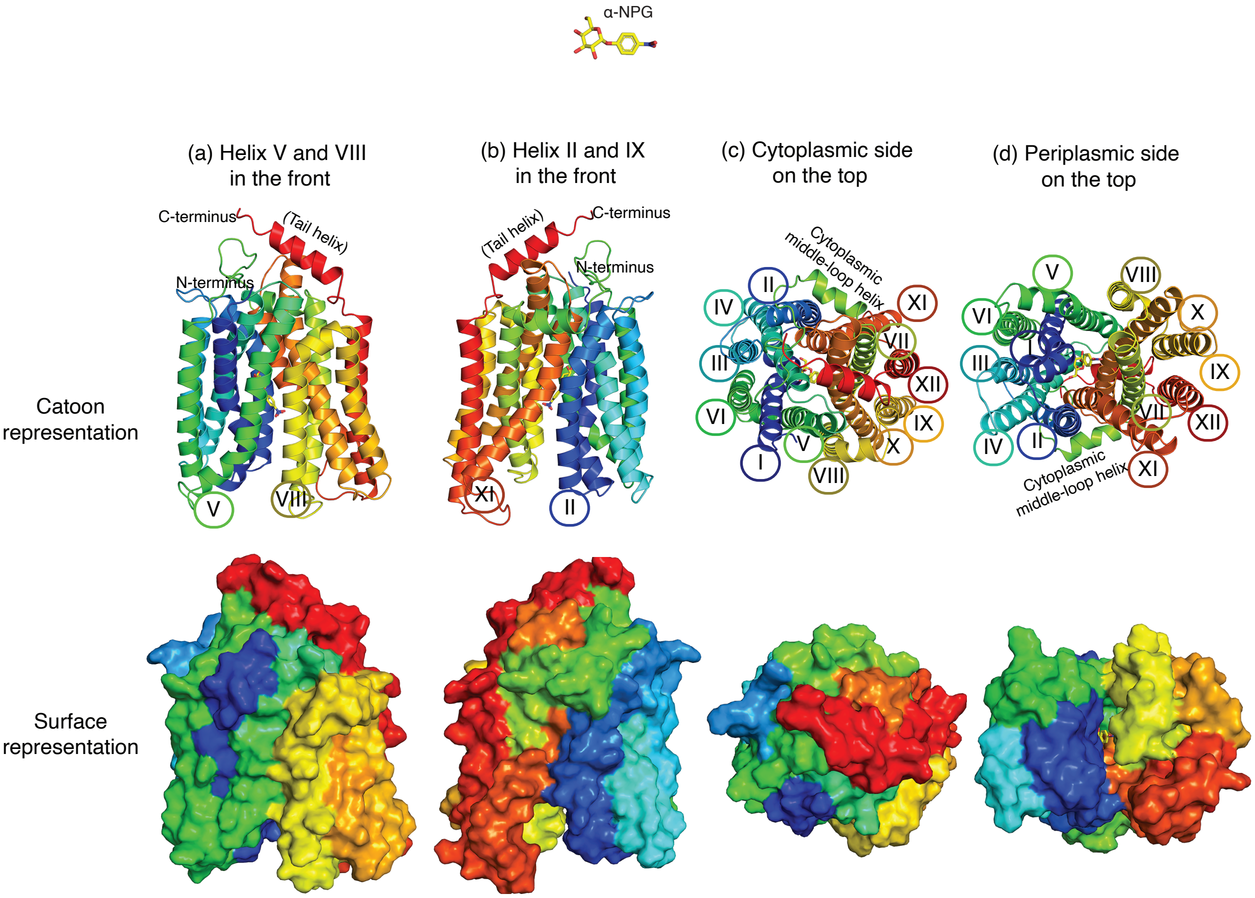
