## Supplementary material for "Molecular recognition of sugar binding in a melibiose transporter MelB by X-ray crystallography": SI Fig. S4

**Supporting Information Fig. S4**

**(a) Cross-eye stereo view of electron density map overlayed on D59C $MelB_{St}$ structure**

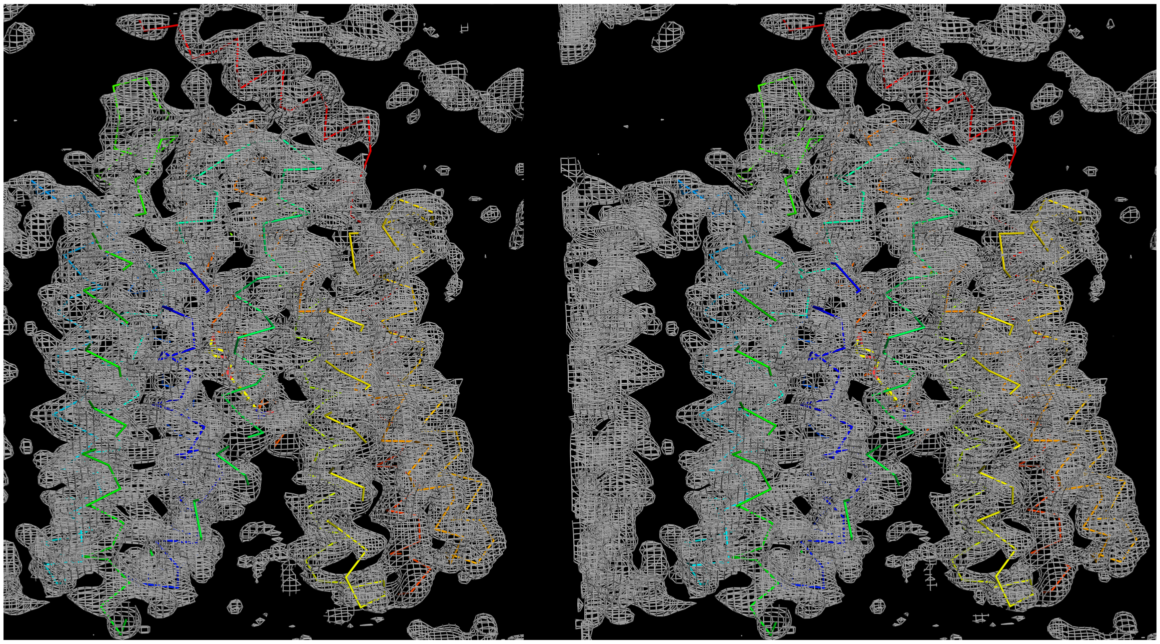

**(b) Cross-eye stereo view of 18 heavy atoms overlayed on D59C $MelB_{St}$ structure**

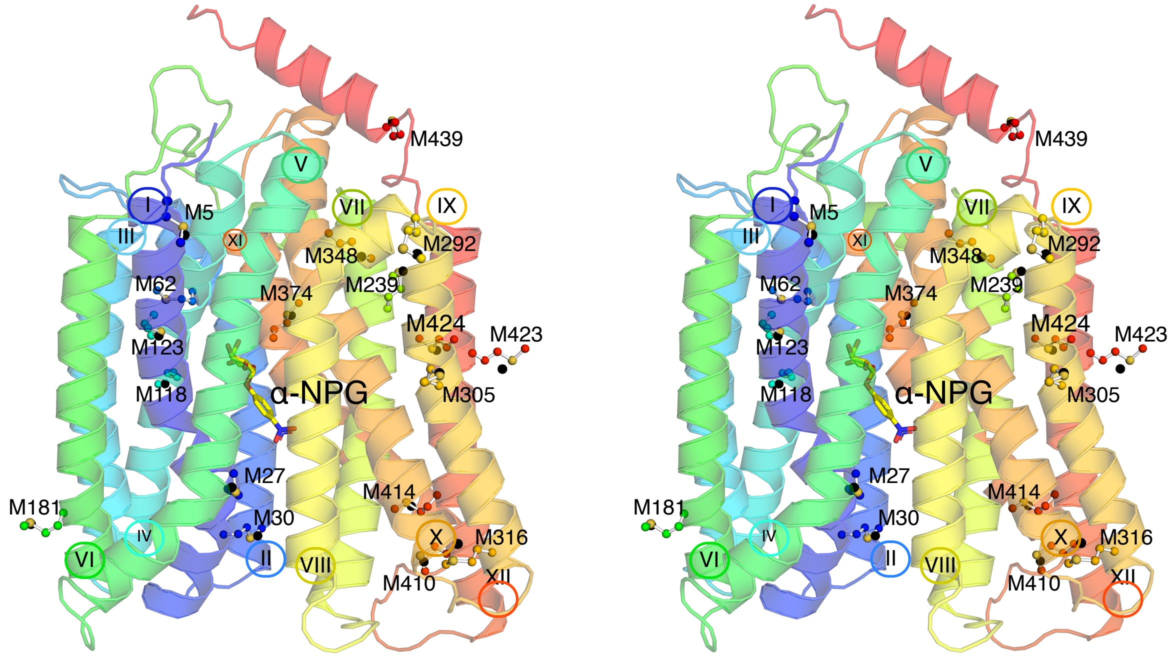
