## Supplementary material for "Molecular recognition of sugar binding in a melibiose transporter MelB by X-ray crystallography": SI Fig. S6

Supporting Information  Fig.S6

### (a) Overview of sugar site and cation site of D59C MelB$_{St}$

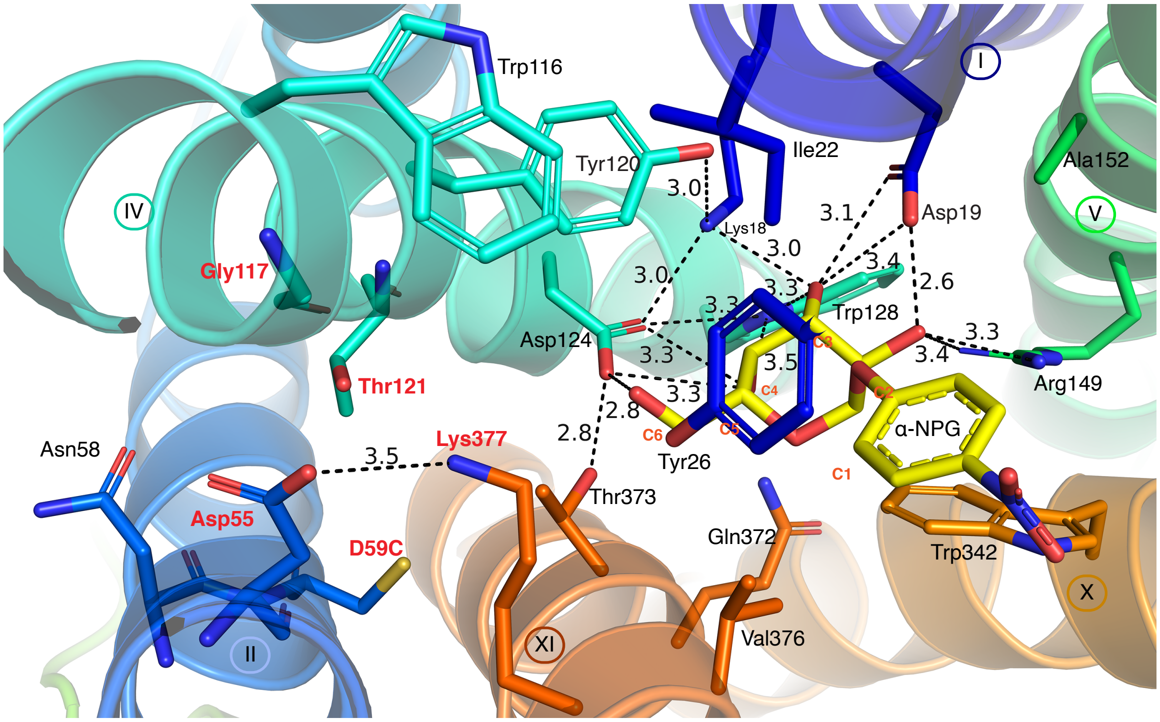


### (b) Multiple sequence alignment focused on the active site

| | | I (18, 19, 22, 26) | II (55, **59**) | IV (117, 120, 121, 124, 128) | V (149) | X (342) | XI (373, 376, 377) |
|---|---|---|---|---|---|---|---|
| P30878 | MELBSt | 18-KDFAIGIVY-26 | 55-DAIND-59 | 117-GMTYTIMDIPFW-128 | 149-R | 342-W | 373-TMVVK-377 |
| Q7VAQ3 | Q7VAQ3_PROMA | 11-TGLAATQLG-19 | 48-DAIND-52 | 110-MTAYTSVNLPFA-121 | 142-F | 336-Y | 371-VLIQK-375 |
| Q8R8R3 | Q8R8R3_THETN | 50-IGIMLEAFG-58 | 87-FAIFN-91 | 148-DFLYVLVGLNLA-159 | 180-Q | 356-V | 387-GFIIR-391 |
| Q65SP4 | Q65SP4_MANSM | 20-CNLVFSTMA-28 | 58-DAVTD-62 | 119-SLAYTVVNIPLN-130 | 151-M | 335-F | 365-GFFVK-369 |
| Q65T75 | MELB_MANSM | 18-KDFAIHIVY-26 | 55-DAVND-59 | 117-GMTYTLMDIPFW-128 | 149-R | 342-W | 374-TMVVK-379 |
| O07366 | MELB_KLEAE | 18-KDFAIGIVY-26 | 55-DAIAD-59 | 117-GFTYTIMDVPFW-128 | 149-R | 342-W | 373-TLVVK-377 |
| Q02581 | MELB_KLEPN | 18-KDFAIGIVY-26 | 55-DAIAD-59 | 117-GFTYTIMDVPFW-128 | 149-R | 342-W | 373-TLVVK-377 |
| P02921 | MELB_ECOLI | 18-KDFAIGIVY-26 | 55-DAIND-59 | 117-GMTYTIMDIPFW-128 | 149-R | 342-W | 373-TMVVK-377 |
| Q83PA7 | MELB_SHIFL | 14-KDFAIGIVY-22 | 51-DAIND-55 | 113-GMTYTIMDIPFW-124 | 149-R | 342-W | 369-TMVVK-373 |
| Q8XDU0 | MELB_ECO57 | 18-KDFAIGIVY-26 | 55-DAIND-59 | 117-GMTYTIMDIPFW-128 | 149-R | 342-W | 373-TMVVK-377 |
| Q9KWG6 | MELB_CITF | 18-KDFAIGIVY-26 | 55-NAFND-59 | 116-GMTYTIMDIPFW-127 | 148-R | 341-W | 372-TMVVK-376 |
| Q8Z1N9 | MELB_SALTI | 18-KDFAIGIVY-26 | 55-DAIND-59 | 117-GMTYTIMDIPFW-128 | 149-R | 342-W | 373-TMVVK-377 |
| Q57GS8 | MELB_SALCH | 18-KDFAIGIVY-26 | 55-DAIND-59 | 117-GMTYTIMDIPFW-128 | 149-R | 342-W | 373-TMVVK-377 |
| Q5PJ19 | MELB_SALPA | 18-KDFAIGIVY-26 | 55-DAIND-59 | 117-GMTYTIMDIPFW-128 | 149-R | 342-W | 373-TMVVK-377 |
| Q14CX5 | MFSD13A_HUMAN | 26----TTILHN-31 | 55-DAIND-59 | 135-GMTYTIMDLHHH-146 | 167-S | 375-C | 408-ALVTK-412 |
| Q8NA29 | MFSD2A_2_HUMAN | 56-YQVTGCALG-64 | 93-DAITD-97 | 155-ETMVTCFHVPYS-166 | 187-M | 403-W | 432-VFFTK-436 |
| A6NFX1 | MFSD2B_HUMAN | 54-NQIASSATA-62 | 91-GAAAD-95 | 153-QALATFFQVPYT-164 | 185-M | 390-W | 419-VFFTK-423 |
